# A genome-scale ORF screen for resilience factors that protect the RPE and restore vision in aged mice

**DOI:** 10.64898/2026.09.29.755187

**Authors:** Yuancheng Ryan Lu, Han Shen, Kathrin Kajderowicz, Jiahe Ai, Yi Hua Chen-Neumann, Yan Hu, Alexander Tyshkovskiy, Qiannuo Lou, Tianwei Dong, Louisa Grauvogel, Zhaoyi Chen, James C. Cameron, Ximeng Li, Shintaro Shirahama, Michael Bligh, Shurong Hou, Jessica Chin, Mauro W. Costa, Jiwon Ryu, Ruchi Sharma, Matthias Mann, Kapil Bharti, Catherine Bowes Rickman, Vadim N. Gladyshev, Meredith Gregory-Ksander, Bruce R. Ksander, Jonathan Weissman

## Abstract

Aging is a complex process, yet therapeutic discovery has largely relied on hypothesis-driven testing of individual targets. Here, we performed an unbiased genome-scale open reading frame (ORF) screen for oxidative-stress resistance in retinal pigment epithelium (RPE) cells, which are particularly vulnerable to oxidative damage with age. The screen identified nine protective ORFs, including antioxidant factors and a regulator of epithelial-to-mesenchymal transition. Notably, we also identified three factors that reduce transcriptomic age, including the cardiac transcription factor NKX2-5. NKX2-5 conferred robust oxidative resilience *in vitro* and *in vivo*. Domain dissection showed that its homeodomain was dispensable for resilience, enabling development of NKX2-5ΔHD, which retained protective activity without detectable toxicity over 14 months. In middle-aged mice, subretinal NKX2-5ΔHD restored visual and electrophysiological function, and its systemic delivery improved grip strength and reduced frailty in late-aged mice. These findings establish genome-scale ORF screening as a strategy to discover transferable resilience factors that can be reengineered to counter age-related functional decline across tissues.

## Introduction

Age is a major risk factor for many chronic and life-threatening diseases, which are complex, multifactorial, and often lack effective therapies^1,2^, and population aging is increasing their global burden. Understanding the molecular basis of aging and natural ways to reverse aging holds the promise of developing transformative interventions that target shared mechanisms of aging, and may help prevent or delay multiple age-related diseases.

At the molecular level, aging is associated with a progressive decline in cellular stress resistance, driven by DNA damage, mitochondrial dysfunction, and impaired protein homeostasis^3^. As these core cellular repair and maintenance systems break down over time, cells lose their resilience and ability to respond to physical, chemical, or biological stress^4,5^.

Oxidative stress arises when the production of reactive oxygen species (ROS) exceeds cellular antioxidant defenses. Young cells generally have the capacity to manage ROS and limit the accumulation of oxidative damage. However, reduced resilience to oxidative stress has been observed during aging in a broad range of age-related diseases^6–8^, including retinal diseases^9^, neurodegenerative disorders^10^, diabetes^11^, chronic lung^12^, kidney^13^ and liver diseases^14^, hearing loss^15^, and cancer^16^. Nevertheless, the endogenous genetic programs that confer oxidative resilience, and how they change during aging, remain incompletely understood, and have not been rigorously interrogated using unbiased genetic screening approaches.

Recent developments in functional genomics have greatly expanded our ability to conduct unbiased systematic forward genetic screens for desired phenotypes. While over-expression screens are more technically demanding and less widely used^17,18^, they offer key advantages over knockout screens, such as a single-step infection and identification of candidates for further gene therapy development^19^. In addition, novel gene functions can be revealed through ectopic expression - a method that led to the seminal discovery of the Yamanaka factors^20^, which we and others have optimized to rejuvenate retinal ganglion cells^21–24^ and other tissues^25–28^, and can extend mouse lifespan when delivered to internal organs^29,30^. With additional data showing efficacy in non-human primates^23^, the OSK (OCT4, SOX2, and KLF4; three Yamanaka factors) gene therapy is now being evaluated in a phase 1 trial (NCT07290244) assessing safety in patients with open-angle glaucoma or non-arteritic anterior ischemic optic neuropathy (NAION), a form of optic neuropathy.

A cell type vulnerable to age-related oxidative stress provides a suitable system for such overexpression screens. The causal role of oxidative stress in disease development is arguably most well-established in ocular diseases, particularly dry age-related macular degeneration (AMD), a leading cause of blindness affecting nearly 200 million people world-wide ^31^, with limited effective therapies^32,33^. Oxidative stress– induced degeneration of retinal pigmented epithelium (RPE) cells is considered a central event in AMD pathogenesis^9,34^. The RPE’s critical location between the highly oxygenated choroid and photo-oxidatively stressed retina makes it highly vulnerable to oxidative stress^35^, such that additional oxidative stress from lifestyle factors and mitochondrial dysfunction during aging can overwhelm its natural defenses. Clinical support for this mechanism comes from the AREDS2 studies, which showed that oral antioxidant supplementation can slow disease progression^36^.

In this study, we performed, to our knowledge, the first genome-scale human open reading frame (ORF) screening to directly select for oxidative-stress resilience in RPE cells. By elucidating novel mechanisms of oxidative defense and nominating genetic interventions, this study seeks to broaden our understanding of cellular resilience, identify therapeutic targets and provide new insights into natural rejuvenative programs.

## Results

### Oxidative injury induces aging-associated features in RPE

To perform a genome-scale search for stress resilience interventions, we needed a scalable cell-culture system. We considered primary RPE, iPSC-derived RPE^37^ and ARPE-19 cells, and chose ARPE-19 because they are readily expandable for a screen requiring ~160 million cells, are more amenable to lentiviral transduction than primary or iPSC-derived RPE, and have not passed through a pluripotent state that could reset molecular age and stress resilience^38^. To mitigate the known limitations of undifferentiated ARPE-19 cells^39,40^, we used an optimized nicotinamide-based maturation protocol to differentiate them at confluence before stress challenges^41,42^, and also used mouse models to validate the top candidates.

We chose sodium iodate (NaIO_3_) as the screening stressor because it induces robust oxidative injury selective for RPE^6,43^, and induces key features associated with RPE aging. In mice, 24 hrs after NaIO_3_ treatment, oxidative stress measured by the CellROX fluorogenic probe increased in mouse RPE (Fig. 1, A to C), but not neuroretina (fig. S1A), to levels comparable to those observed in aged animals, indicating that this injury paradigm acutely elevates oxidative burden in RPE cells *in vivo*. In ARPE-19 cells, NaIO_3_ activated aging-associated inflammatory, interferon, EMT and metabolic-stress programs (Fig. 1, D and E). Thus, NaIO_3_ provides a scalable oxidative-stress paradigm to identify genes that enhance RPE resilience, as used in our prior study^44^.

**Fig. 1.**
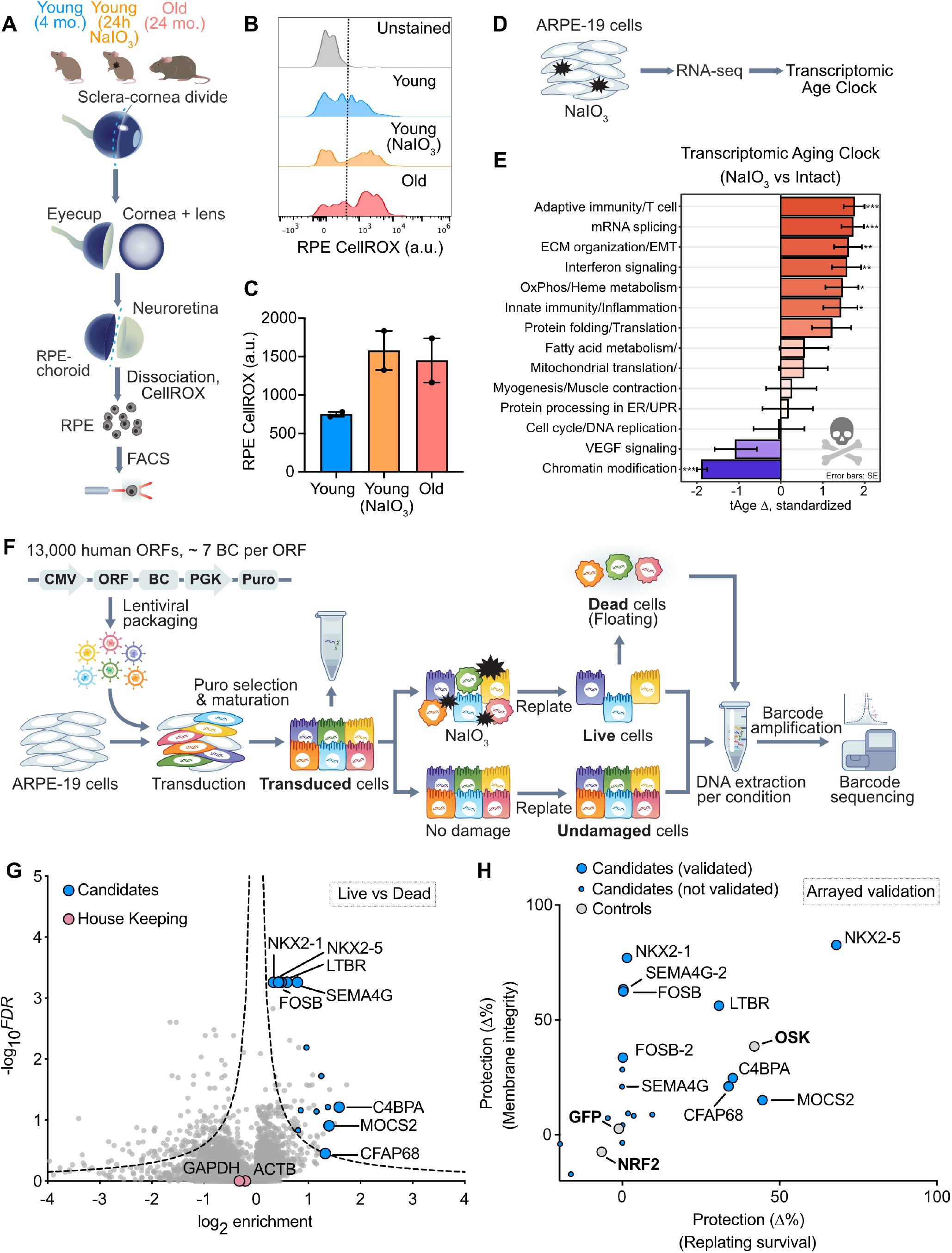
Aging and NaIO_3_-induced injury increase oxidative stress in the RPE, enabling an ORF screen for protective factors. **(A)** Experimental workflow for isolating RPE cells from young (4-month-old), NaIO_3_-treated young, and aged (24-month-old) mice for ROS quantification by CellROX staining and flow cytometry. **(B)** Representative flow cytometry histograms showing increased CellROX DeepRed signal in live RPE cells from NaIO_3_-treated young mice and aged mice compared with young controls. **(C)** Quantification of CellROX fluorescence intensity from live RPE cells in young, NaIO_3_-treated young, and aged mice (n=2 in each group). Data are shown as mean ± s.e.m. **(D)** Schematic of NaIO_3_ treatment in ARPE-19 cells followed by RNA-seq and transcriptomic age-clock analysis. **(E)** Module-specific transcriptomic aging-clock analysis of ARPE-19 cells after NaIO_3_ treatment, showing induction of aging-associated inflammatory, immune, extracellular matrix, metabolic, and proteostasis-related modules. **(F)** Schematic of the pooled human ORF screen. ARPE-19 cells were transduced with a barcoded ORF library, selected and matured, challenged with NaIO_3_ or left untreated, and live, dead, and undamaged populations were collected for barcode sequencing. **(G)** Volcano plot showing differential ORF enrichment in live versus dead cells after NaIO_3_ challenge. Each dot represents a single ORF. Gray dots indicate all screened ORFs, pink dots denote housekeeping genes, and blue dots highlight selected candidate ORFs subsequently tested in the arrayed validation assay. Larger/smaller blue dots indicate validated/non-validated hits respectively. **(H)** Arrayed validation of candidate protective factors after oxidative challenge,Δ%, + DOX relative to - DOX condition. Protection was quantified using two complementary readouts: replating survival and membrane integrity. Validated candidates are shown as large blue circles, non-validated candidates as small blue circles, and positive and negative controls as grey circles.

### Genome-scale ORF screen in RPE cells

We used a lentiviral library of human ORFs that contained nearly 13,000 full-length ORFs^17^, with approximately 7 barcodes per ORF. We amplified the library, produced lentivirus, and transduced 160 million ARPE-19 cells at a 35% infection rate, resulting in over 4,000x representation per ORF after puromycin selection of transduced cells. The transduced ARPE-19 cells captured 99.8% of all ORFs, including the larger ones (fig. S1B) and ORF representation was highly correlated between technical replicates and with the plasmid library (fig. S1, C and D). Transduced ARPE-19 cells were challenged with NaIO_3_ for 18h ^45^ and then replated into new culture dishes with growth medium. ARPE-19 cells that survived the NaIO_3_ treatment and reattached in the replated cultures were harvested and sequenced to identify their ORF barcodes. As internal controls, the floating dead cells from these cultures were also harvested and their ORF barcodes sequenced (Fig. 1F). Finally, the ORF barcodes were obtained from undamaged, replated ARPE-19 cells as an additional control for effects on proliferation (fig. S1E). In all these conditions, over 99.6% of all potential ORFs were captured for analysis, indicating a robust screen that provided genome-scale coverage of candidate genes (fig. S1F). Candidate protective ORFs that promoted resilience were ranked by barcode enrichment in surviving cells. ORFs with multiple enriched barcodes generally showed higher significance, but highly enriched ORFs with fewer barcodes were also included for downstream arrayed validation (fig. S1G).

As a control for nonspecific growth effects, we asked whether the prioritized hits simply conferred a proliferative advantage, which could raise potential safety concerns if proliferation was induced *in vivo*. We therefore compared ORF abundance in untreated cells after replating and expansion for 4 days with the transduced baseline population (fig. S1H). Established growth-promoting factors, including *MYC, LIN28A* and *WRAP53*, were enriched after expansion, whereas growth-inhibitory genes, including *RASSF1, CABLES1, IGFBP5* and *PTPRO*, were depleted, demonstrating that the screen was capable of detecting ORFs that alter cellular fitness. By contrast, the prioritized ORF candidates were not enriched during the 4-day expansion, suggesting that their selection was unlikely to be driven by a general proliferative advantage under these conditions.

Seventeen prioritized ORF candidates and their isoforms were each cloned into inducible lentiviral vectors to transduce ARPE-19 cells, and both membrane integrity and replating survival assays were used to assess their protection against oxidative injury following NaIO_3_ treatment (Fig. 1G and fig. S1I). GFP and OSK served as negative and positive controls, respectively (Fig. 1H).

Nine ORF candidates: *NKX2-5, LTBR, C4BPA, CFAP68, MOCS2, NKX2-1, SEMA4G*, and two isoforms of *FOSB* exhibited strong protection in arrayed validation (Fig. 1H and fig. S1, J and K). These candidate genes show varied expression patterns across RPE aging and disease as assessed in Spectacle^46^, a scRNA-seq database from human donor ocular samples (fig. S2A). For example, FOSB expression is higher in infant RPE cells but also significantly upregulated in neovascular AMD, and therefore may warrant additional evaluation before therapeutic use. In addition, outside the eye, these ORFs are expressed at varying levels across different tissues (fig. S2B). Interestingly, *NKX2-5* and *NKX2-1* were not detectably expressed in the RPE datasets examined, but are highly expressed in the heart and lung, respectively— two organs exposed to high oxygen tension. Thus, ectopic expression of ORFs in cell types where they are not normally expressed can reveal unexpected functions.

### Protective ORFs induce distinct transcriptional programs

To assess the similarities and functional differences among the nine validated protective ORFs, we conducted RNA-seq after 4 days of individual ORF expression in ARPE-19 cells.

*NRF2* and *OSK* expression and TNFα treatment were included as comparators. By calculating the correlation of pathway enrichment profiles, we classified these nine ORFs into three distinct classes based on their transcriptome signatures (Fig. 2A).

**Fig. 2.**
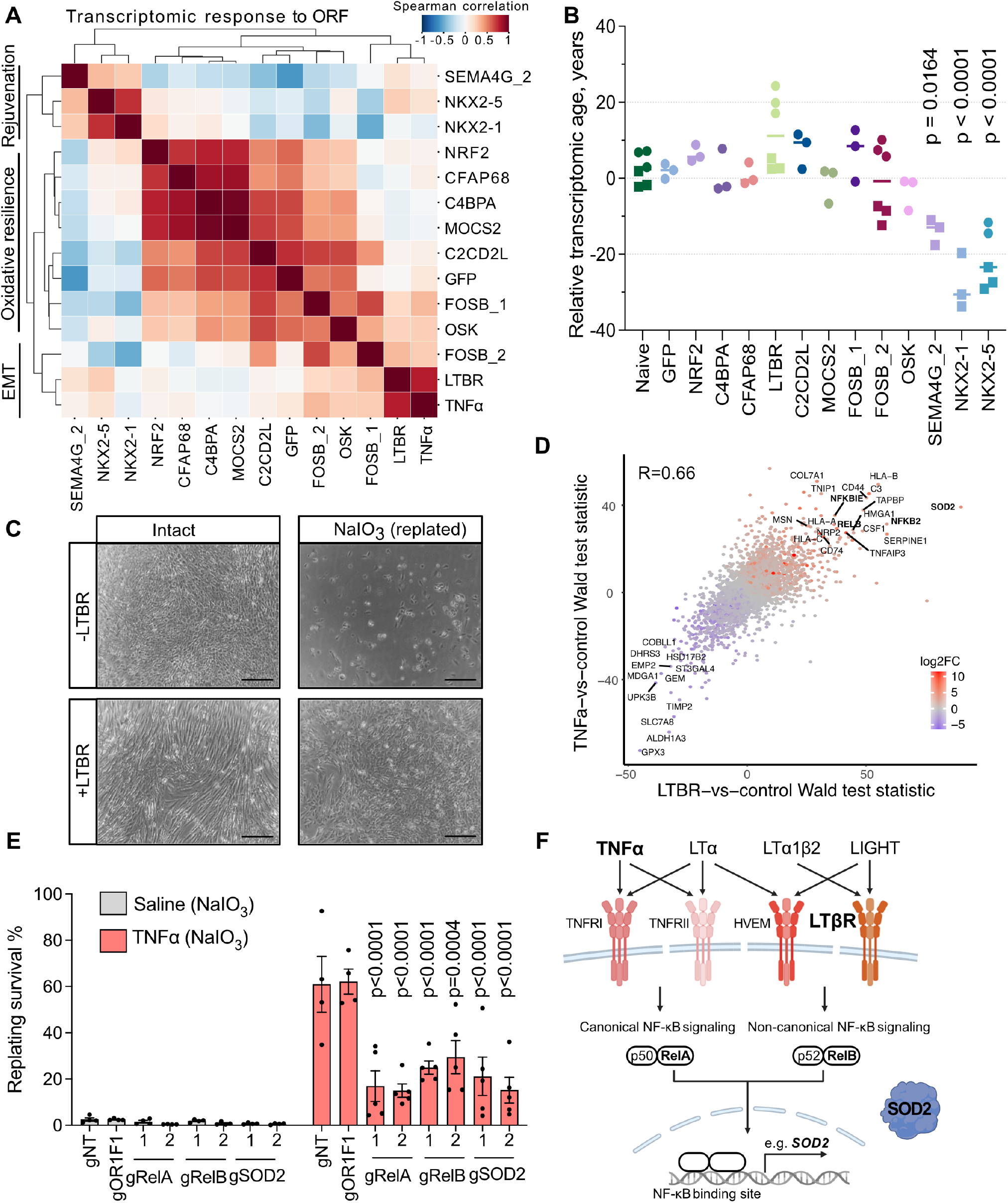
Transcriptional comparison of protective ORFs identifies distinct oxidative-resilience, EMT, and rejuvenation-associated programs. **(A)** Spearman correlation matrix calculated using GSEA enrichment of differential gene expression induced by protective ORF overexpression in ARPE-19 cells, showing relationships among oxidative-resilience-, EMT-, and rejuvenation-associated candidates. All normalized to batch matched naive untreated controls. **(B)** Relative transcriptomic age estimates of ARPE-19 cells after ORF overexpression, calculated with the multi-species multi-tissue chronological clock. Each dot represents an independent sample. P values were calculated by comparison with naive untreated ARPE-19 using one-way ANOVA. **(C)** Representative bright-field images of ARPE-19 cells with doxycycline-inducible LTBR expression, with or without NaIO_3_ challenge. Scale bar, 50μm. **(D)** Correlation of genome-wide differential expression signatures induced by LTBR overexpression and TNFα treatment, shown as Wald test statistics relative to control (naive untreated control for TNFα, GFP control for LTBR). Using the GFP control for LTBR accounts for the effects of DOX induction; however, the correlation was similar (r = 0.67) when using batch-matched naive controls. Selected shared upregulated and downregulated genes are labeled. **(E)** Replating survival of TNFα-treated ARPE-19 cells after NaIO_3_ challenge following CRISPRcut targeting of *RelA, RelB*, or *SOD2*. Non-targeting guide (gNT) and *OR1F1*-targeting guide (gOR1F1) are shown as controls. P values were calculated by two-way ANOVA with comparisons against the gNT control. **(F)** Schematic of TNF-superfamily ligand–receptor signaling through canonical and non-canonical NF-κB pathways, including downstream induction of candidate antioxidant effectors such as SOD2.

One class, including *MOCS2, C4BPA*, and *CFAP68*, as well as *FOSB*, which clustered closely with *OSK*^*44*^, grouped near *NRF2*, suggesting related antioxidant transcriptional programs. A second class was exemplified by LTBR, a receptor in the TNF superfamily, while the third comprised *NKX2-1, NKX2-5*, and *SEMA4G* (Fig. 2A). Using a recently established transcriptomic aging clock^47^ that distinguishes young from old mouse RPE cells (fig. S2C), we tested whether these factors have an effect on molecular age. Notably, *NKX2-1, NKX2-5* and *SEMA4G* overexpression shifted transcriptomic age estimates toward younger values (> 15 years) (Fig. 2B), suggesting their effects on age-associated transcriptional programs.

We were first intrigued by LTBR-mediated oxidativestress resistance, as *LTBR* overexpression induced an EMT-like morphology (Fig. 2C) and a transcriptomic signature resembling those observed in TNFα-treated RPE cells (Fig. 2D). However, TNF*α* has been previously associated with AMD pathogenesis, not protection. Increased levels of TNF*α* were detected in the vitreous^48^ and serum^49^ of AMD patients and TNF*α* is known to upregulate *RelA* and *RelB*, which activate NF-κB-driven EMT, a phenomenon frequently observed with RPE dysfunction in geographic atrophy and at the fibrotic end stage of neovascular AMD^50,51^(Fig. 2D).

To confirm it is not due to a non-canonical function of LTBR, we tested four TNF-superfamily ligands—TNFα, LTα, LTα1β2, and LIGHT—and found that all conferred protection against oxidative injury to varying degrees (fig. S2D). Mechanistically, we observed that *SOD2*, a downstream antioxidant gene responsive to NF-κB signaling, was strongly upregulated by both *LTBR* overexpression and TNFα. Using CRISPR-Cas9 to knock out *RelA, RelB*, and *SOD2*, we found that the oxidative protection conferred by TNFα was abolished. In contrast, knockout of *OR1F1*, a control gene not expressed in RPE cells, had no effect (Fig. 2E and fig. S2E). The failure of *SOD2* to emerge as an ORF-screen hit can be explained by the presence of two SOD2 isoforms in the library: the full-length isoform was significantly protective, whereas the truncated isoform conferred no protection and likely diluted the overall signal (fig. S2F).

Overall, these data support a requirement for the NF-κB/SOD2 axis in TNFα-mediated protection against oxidative injury in RPE cells (Fig. 2F), and except *LTBR*, all other validated candidates do not upregulate this axis (fig. S2G), suggesting that the TNF super family signaling may function as an endogenous resilience pathway.

### NKX2-5 protects RPE cells and improves visual acuity in aged mice

Among the three candidate factors demonstrating the ability to reduce transcriptomic age, we focused on NKX2-5 not only because it showed the strongest protection against NaIO_3_-induced oxidative stress *in vitro* (Fig. 1H and Fig. 3A) but also because of its endogenous expression pattern, which is largely restricted to the heart, while being silenced in most other tissues (Fig. 3B, fig. S2B, and fig. S3A).

**Fig. 3.**
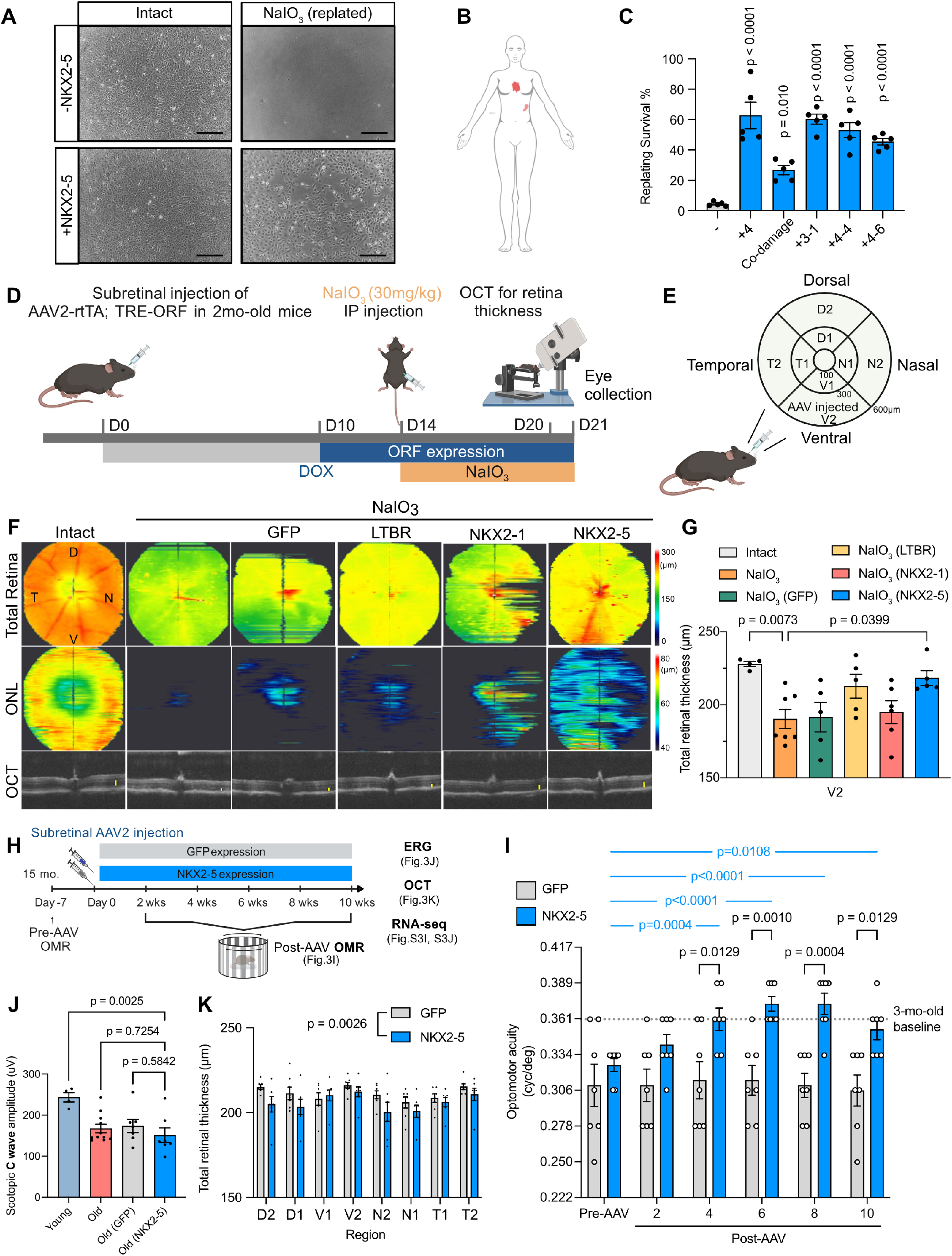
NKX2-5 protects against oxidative retinal injury and improves visual acuity in aged mice but causes toxicity with prolonged expression. **(A)** Representative bright-field images of ARPE-19 cells with or without NKX2-5 expression under intact conditions or after NaIO_3_ challenge followed by replating. Scale bar, 50μm. Quantification is shown in **Fig.S1K and 3C**. **(B)** Schematic indicating endogenous NKX2-5 expression in the heart. **(C)** Replating survival of ARPE-19 cells expressing NKX2-5 under the indicated treatment regimens after NaIO_3_ challenge. **(D)** Experimental timeline for testing AAV-mediated ORF expression in an acute NaIO_3_-induced RPE injury model. Two-month-old mice received subretinal AAV-EF1α-rtTA; TRE-ORF injection, followed by doxycycline induction, systemic NaIO_3_ administration (30 mg kg^−1^), OCT imaging, and tissue collection. **(E)** Schematic of retinal subregions used for OCT-based thickness analysis, defined by radial distance from the optic nerve head and retinal orientation. D, dorsal; V, ventral; N, nasal; T, temporal. **(F)** En face retinal thickness maps, ONL thickness maps, and representative OCT cross-sectional images from intact and NaIO_3_-treated eyes expressing GFP, LTBR, NKX2-1, or NKX2-5. Yellow lines indicate ONL thickness measurements in OCT images. **(G)** Quantification of total retinal thickness in the untreated eye (n=4), NaIO_3_ injured eye (n=7), AAV-injected ventral region V2 after NaIO_3_ injury in eyes expressing GFP, LTBR, NKX2-1, or NKX2-5 (n=5, 5, 6 and 5 respectively). **(H)** Experimental timeline for assessing AAV-mediated GFP or NKX2-5 expression in aged mice. Fifteen-month-old mice received subretinal AAV2-GFP or AAV2-NKX2-5 injection after baseline OMR testing, followed by longitudinal OMR analysis and endpoint ERG, OCT, RNA-seq analyses. **(I)** Longitudinal OMR measurements of visual acuity in aged mice (15-month-old) after subretinal AAV-GFP (n=7) or AAV-NKX2-5 (n=7) injection. The dotted line indicates the 3-month-old baseline reference. Two-way repeated-measures ANOVA with Šídák multiple comparisons between groups at each time point and versus pre-AAV baseline within groups. Examiners were masked to group allocation. **(J)** Scotopic ERG c-wave amplitudes in young mice (n=4), untreated old mice (n=11), and old mice 10 weeks after AAV-GFP (n=7) or AAV-NKX2-5 (n=7) injection. **(K)** Regional OCT quantification of total retinal thickness in aged (15-month-old) mice 10 weeks after AAV-GFP (n=7) or AAV-NKX2-5 (n=7) injection. D, dorsal; V, ventral; N, nasal; T, temporal.

We found that protection conferred by NKX2-5 was both rapid and durable: transient NKX2-5 expression remained protective even 6 days after NKX2-5 protein was no longer detected (Fig. 3C and fig. S3, B and C), suggesting a durable effect. NKX2-5 also protected against complement-mediated injury, another AMD-associated stressor^52^ (fig. S3D). Although NKX2-5 is canonically known for its developmental role in cardiogenesis through interactions with GATA4, TBX5 and MYH6, this cardiac gene network was not induced in NKX2-5-expressing ARPE-19 cells (fig. S3E), consistent with the absence of overt morphological changes (Fig. 3A). These findings suggest that NKX2-5 has a previously unrecognized function in oxidative resilience that is separable from its canonical cardiogenic activity.

To test whether NKX2-5 protects the RPE and retina *in vivo*, we used the acute NaIO_3_-induced RPE degeneration model^53^ and generated all-in-one doxycycline-inducible AAV2s expressing either GFP, LTBR, NKX2-1 or NKX2-5. Two-month-old mice received subretinal AAV2 injection, followed by 4 days of ORF induction before systemic NaIO_3_ challenge for 7 days (Fig. 3D). As expected, systemic NaIO_3_ caused RPE injury and secondary loss of total retinal and outer nuclear layer thickness (Fig. 3, E to G). Interestingly, OCT analysis near the injected region showed that NKX2-5 markedly preserved total retinal and outer nuclear layer thickness after injury, outperforming GFP, LTBR and NKX2-1 (Fig. 3, E to G, and fig. S3F). These findings identify NKX2-5 as a potent *in vivo* protector against acute RPE degeneration.

Because age-related loss of oxidative resilience and ROS accumulation contribute to RPE degeneration, and aging is a primary risk factor for AMD^54^, we next tested whether NKX2-5 could restore oxidative resilience in aged RPE. Because AAV2 vectors with a constitutive CAG promoter efficiently target RPE following subretinal injection, an approach used in an FDA-approved gene therapy^55,56^, we injected AAV2-CAG-GFP and AAV2-CAG-NKX2-5 into the subretinal space of 15-month-old mice (Fig. 3H). We tracked their visual acuity using the OptoDrum instrument before and after AAV injection. Longitudinal OptoDrum analyses showed no improvement in GFP-treated eyes, whereas NKX2-5-treated eyes showed significantly improved visual acuity starting after 4 weeks of treatment, approaching the mean values in young (Fig. 3I).

Visual acuity in NKX2-5-treated eyes remained significantly above the pre-AAV baseline, although the benefit began to diminish after 8 weeks. By 10 weeks, OCT revealed a significant reduction in retinal thickness, accompanied by a nonsignificant decline in ERG responses (Fig. 3, J and K, and fig. S3, G and H). This type of discrepancy between OMR results and ERG results has been reported previously^57^ and by us with AAV2-OSK^44^ (Supplementary Discussion). Transcriptomic age (tAge) analysis^47^ provided some molecular context for these findings. Although ectopic NKX2-5 induced youthful mitochondrial and inflammatory programs, it also activated aging-associated pathways, including mTOR signaling and mitochondrial translation/respiration (fig. S3I). Collectively, these changes increased the transcriptomic age of NKX2-5-treated RPE (fig. S3J), similar to our previous findings that OSK expression in RPE cells increased transcriptomic age despite concurrent improvements in OMR-measured visual acuity^44^. Together, these findings suggest that prolonged NKX2-5 expression engages both protective and detrimental molecular pathways in aged RPE cells, motivating identification of the domain responsible for resilience so that the beneficial effects can be retained while minimizing toxicity.

### Homeodomain deletion preserves NKX2-5-mediated oxidative protection and eliminates toxicity

NKX2-5 protein has four recognized domains: the Tinman domain (TN), homeodomain (HD), NK-2-specific domain (NK-2 SD), and tyrosine-rich domain (YRD). Among them, the HD is the best studied and mediates DNA binding and activation of target genes during cardiogenesis^58^(Fig. 4A). *NKX2-5* transgenic mice carrying a germline HD deletion or mutation (I184M or I184P) have reduced DNA binding ability to its canonical targets ^59^ and abnormal heart development, resulting in early death at E10. *NKX2-5* transgenic mice with an inducible systemic or ventricle-targeted HD mutation developed abnormal heart function following induction of the mutation in adults, resulting in early mortality^60^.

**Fig. 4.**
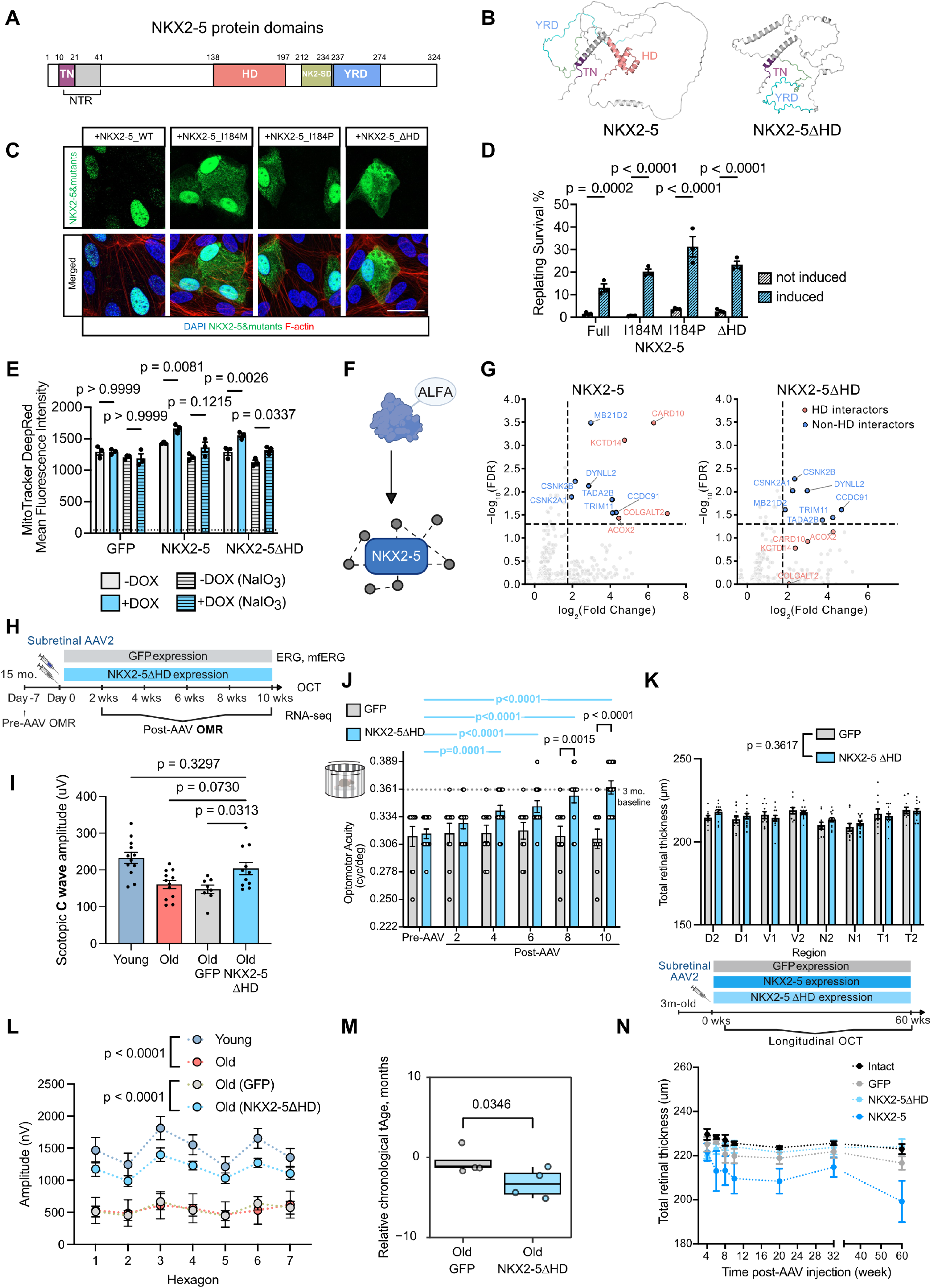
AAV-NKX2-5ΔHD reduces retinal toxicity and improves visual function in aged mice. **(A)** Schematic of NKX2-5 protein domains, including the Tinman domain (TN), homeodomain (HD), NK-2-specific domain (NK-2 SD), and YRD. Another N-terminal structured region (NTR, aa 14–41) is predicted by AlphaFold 3. **(B)** AlphaFold-predicted structures of full-length NKX2-5 and NKX2-5ΔHD. (**C**) Representative immunofluorescence images of ARPE-19 cells expressing full-length NKX2-5, I184M, I184P, or NKX2-5ΔHD. NKX2-5 is shown in green, DAPI staining in blue, and F-actin in red. (**D**) Replating survival of cells expressing full-length NKX2-5 or the indicated NKX2-5 variants, with or without induction. (**E**) Quantification of MitoTracker Deep Red mean fluorescence intensity in cells expressing GFP, NKX2-5, or NKX2-5ΔHD, with or without doxycy-cline-induced transgene expression and NaIO_3_-induced oxidative stress. Data are shown as mean ± s.e.m.; each point represents an independent replicate; dotted line represents unstained values. Statistical significance was determined by the indicated comparisons. (**F**) Schematic of ALFA-tagged NKX2-5 immunoprecipitation strategy used to identify proteins associated with NKX2-5 and its mutants. (**G**) Co-IP–mass spectrometry analysis of ALFA-tagged NKX2-5 and NKX2-5ΔHD binding partners. Volcano plots show enriched interactors for fulllength NKX2-5 and NKX2-5ΔHD. (**H)** Experimental timeline for assessing AAV2-mediated GFP or NKX2-5ΔHD expression in aged mice. Fifteen-month-old mice received subretinal AAV2-GFP or AAV2-NKX2-5ΔHD injection after baseline OMR testing, followed by longitudinal OMR analysis and endpoint ERG, OCT, RNA-seq analyses. (**I**) Longitudinal OMR measurements of visual acuity in aged mice after subretinal AAV-GFP (n=10) or AAV-NKX2-5ΔHD (n=13) injection. Similar results were obtained in two independent experiments and combined for analysis; one experiment included a later time point shown in **Fig. S4B**. The dotted line indicates the 3-month-old baseline reference. Two-way repeated-measures ANOVA with Šídák multiple comparisons between groups at each time point and versus pre-AAV baseline within groups. Examiners were masked to group allocation. (**J**) Regional OCT quantification of total retinal thickness in aged mice 10 weeks after AAV-GFP (n=10) or AAV-NKX2-5ΔHD (n=13) injection. D, dorsal; V, ventral; N, nasal; T, temporal. (**K**)Scotopic ERG c-wave amplitudes in young mice (n=12), untreated old mice (n=12), and old mice 10 weeks post AAV-GFP (n=8) or AAV-NKX2-5ΔHD (n=11), two mice did not remain asleep for testing in each AAV group. (**L**) Multifocal ERG responses across retinal hexagons in young mice (n=8), untreated old mice (n=5), and old mice 10 weeks after AAV-GFP (n=10) or AAV-NKX2-5ΔHD (n=13) injection. (**M**) Relative transcriptomic age estimates of old RPE treated with AAV-GFP (n=4) or AAV-NKX2-5ΔHD (n=4), calculated with the chronological multitissue clock. (**N**) Longitudinal OCT analysis of total retinal thickness at the AAV injection site after AAV-GFP, AAV-NKX2-5, or AAV-NKX2-5ΔHD injection (n=5, 6, 5, 4 eyes respectively). Data are shown as mean ± s.e.m.

However, we hypothesized that NKX2-5-mediated oxidative protection is a noncanonical function aside from cardiogenesis, and that the toxicity after prolonged expression may trigger by HD-mediated activity. Supporting this hypothesis, NKX2-5 with an HD deletion (ΔHD) still has a stable structure based on the AlphaFold structure prediction and therefore it is possible to produce a stable NKX2-5ΔHD protein (Fig. 4B). To test this hypothesis, we generated ARPE-19 cells expressing the NKX2-5 HD point mutants I184M or I184P, or the ΔHD deletion mutant (Fig. 4C) and tested their response to NaIO_3_. Surprisingly, all mutants showed significant protection against NaIO_3_-induced oxidative injury (Fig. 4D). Protection was not associated with reduced ROS levels in this assay: we detected no reduction of ROS when overexpressing either NKX2-5 or NKX2-5ΔHD (fig. S4A), but both treatments preserve the MitoTracker Deep Red signal under oxidative challenge (Fig. 4E).

To exclude the possibility that NKX2-5ΔHD acquires novel protein-protein interactions, we performed affinity-enrichment mass spectrometry (AE-MS) using ALFA-tagged^61^ full-length NKX2-5 and NKX2-5ΔHD (Fig. 4F). All interactors detected with NKX2-5ΔHD were also found with full-length NKX2-5, indicating that HD deletion did not create new interactions but instead altered a subset of existing ones. Several proteins, including KCTD14 and CARD10, were preferentially associated with the homeodomain-containing protein. CARD10 was particularly notable because it was strongly enriched with full-length NKX2-5 but markedly reduced after HD deletion (Fig. 4G). CARD10, also known as CARMA3, is a scaffold protein in signaling complexes that promote NF-κB activation in non-immune cells^62,63^.

Together, these AE-MS findings suggest that the NKX2-5 homeodomain engages a distinct interaction module, with CARD10 providing a plausible link between full-length NKX2-5 and inflammatory or stress-associated toxicity. Homeodomain deletion may therefore improve NKX2-5 safety by preserving oxidative-resilience activity while uncoupling HD-associated interactions that contribute to toxicity.

### AAV-NKX2-5ΔHD sustainably improves visual function in aged mice with long term safety

Motivated by these *in vitro* findings, we next asked whether NKX2-5ΔHD could preserve the beneficial effects of fulllength NKX2-5 while avoiding its toxicity *in vivo*. We delivered AAV2-CAG-GFP or AAV2-CAG-NKX2-5ΔHD into the subretinal space of 15-month-old mice (Fig. 4H), longitudinally assessed visual acuity using the OptoDrum assay, and measured electrophysiology and retinal thickness after 10 weeks. As expected, GFP-treated control eyes again showed no improvement. Excitedly, NKX2-5ΔHD-treated eyes exhibited a significant increase in visual acuity beginning 4 weeks after injection, reaching levels comparable to those of 3-month-old mice by 10 weeks (Fig. 4I). Notably, this improvement was maintained for another 6 months, when mice had reached 21 months of age (fig. S4B).

Unlike AAV2-NKX2-5, AAV2-NKX2-5ΔHD did not reduce retinal thickness at 10 weeks (Fig. 4J). Moreover, NKX2-5ΔHD significantly improved c-wave amplitudes, together with increases in a- and b-wave amplitudes, in contrast to the trend of decline observed with AAV2-NKX2-5 (Fig. 4K, and fig. S4, C and D). Consistent with this, mfERG revealed enhanced light-evoked retinal function in NKX2-5ΔHD-overexpressing eyes compared with untreated or GFP-treated eyes (Fig. 4L and fig. S4E). At the molecular level, NKX2-5ΔHD overexpression also significantly reduced the transcriptomic age of RPE cells (Fig. 4M), as assessed by tAge^47^, which was accompanied by increased expression of genes involved in DNA repair and mitochondrial function and reduced inflammatory and mTOR signaling (fig. S4F). Together, these results indicate that NKX2-5ΔHD improves multiple measures of retinal function and age-associated transcriptional state.

To further evaluate long-term safety, we performed longitudinal OCT imaging in young healthy mice, which better tolerate repeated anesthesia, following subretinal delivery of AAV2-CAG-GFP, AAV2-CAG-NKX2-5, AAV2-CAG-NKX2-5ΔHD^64^. Consistent with our aged-mouse findings, fulllength NKX2-5 caused progressive retinal thinning. In contrast, NKX2-5ΔHD showed no detectable retinal thinning or structural toxicity after as long as 14 months of overexpression (Fig. 4N and fig. S4G), supporting an improved safety profile with continuous expression.

### Transcriptomic and protein-interaction analyses link NKX2-5ΔHD to suppression of hypoxia, inflammation, and stress responses

To identify potential protective regions outside the NKX2-5 homeodomain, we generated, in addition to ΔHD, a series of deletion mutants lacking the TN domain (ΔTN), NK2-specific domain (ΔNK2-SD), tyrosine-rich domain (ΔYRD), as well as a tyrosine-to-alanine YRD mutant (YRD^Y-A^)^65^ and an N-terminal truncation mutant lacking both the homeodomain and C terminus (N1-137). We then investigated their protective activity, localization, transcriptomic responses, and protein interactions (Fig. 5, A and B, and fig. S5, A and B). ΔHD, ΔTN, or ΔNK2-SD mutants preserved oxidative protection activity, ΔYRD or YRD^Y-A^ mutants show reduced protection in survival and membrane-integrity assays. In contrast, N1-137 completely abolished protection (Fig. 5, C and D). Immunostaining showed that full-length NKX2-5 and mutants retaining the homeodomain localized predominantly to the nucleus, whereas ΔHD and N1-137, both lacking the homeodomain, displayed mixed nuclear and cytoplasmic localization (Fig. 5E). These findings are consistent with prior studies that identified basic nuclear-localization motifs within the NKX2-5 homeodomain^66,67^. Together, these functional and localization analyses suggest that non-homeodomain regions of NKX2-5 mediate its protective activity, with the YRD domain as a partial contributor.

**Fig. 5.**
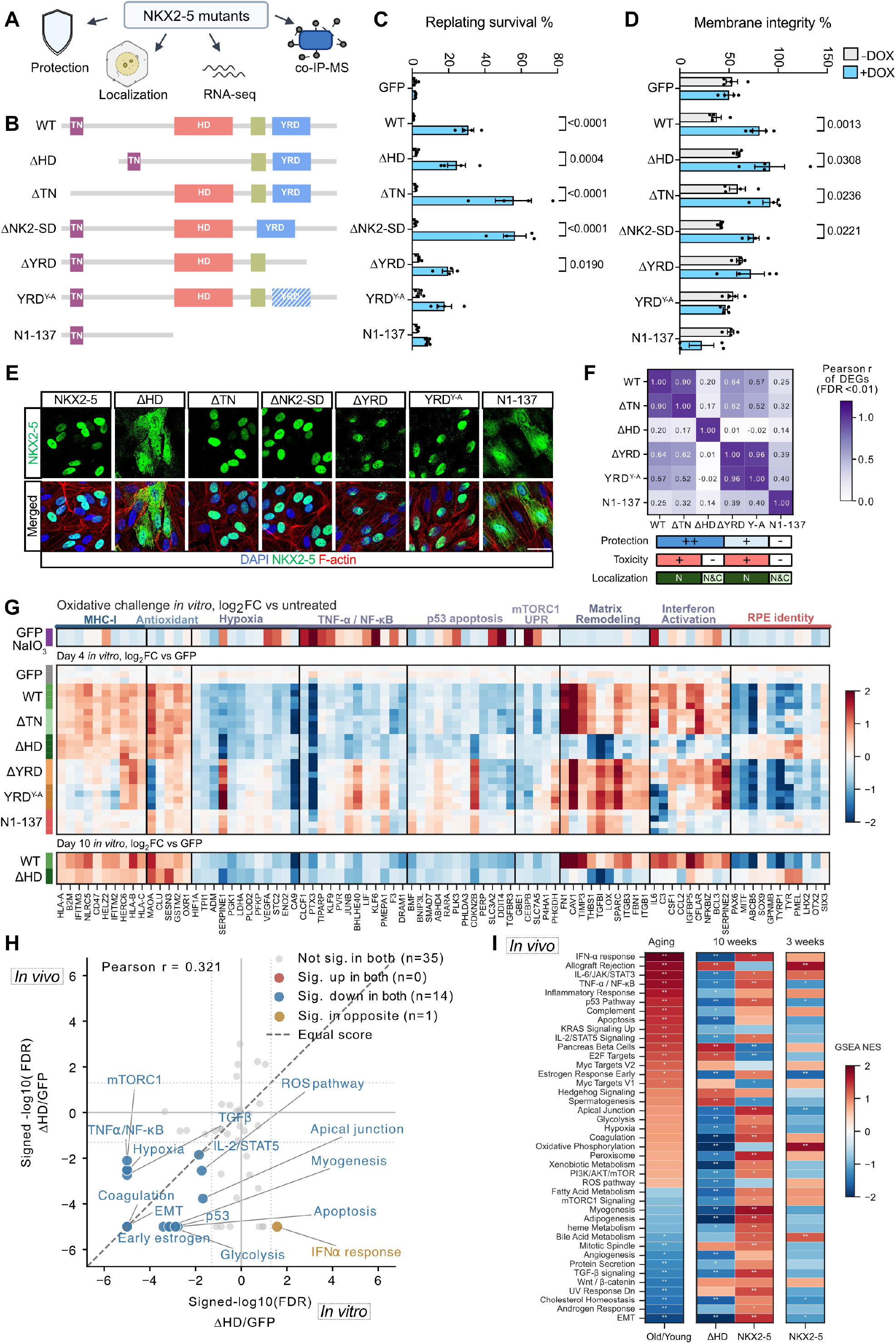
NKX2-5ΔHD induces conserved oxidative-resilience and youth-associated transcriptional programs *in vitro* and *in vivo*. (**A**) Experimental strategy for functional dissection of NKX2-5 mutants by oxidative-stress protection, subcellular localization, RNA-seq, and co-IP– mass spectrometry. (**B**) Schematic of full-length NKX2-5 (WT) and indicated deletion or point-mutant constructs. TN, Tinman domain; HD, homeodomain; YRD, tyrosinerich domain. (**C**) Replating survival of ARPE-19 cells expressing GFP, full-length NKX2-5 or indicated NKX2-5 mutants after NaIO_3_ challenge, with or without doxycycline induction. (**D**) Membrane-integrity assay of ARPE-19 cells expressing GFP, full-length NKX2-5 or indicated NKX2-5 mutants after NaIO_3_ challenge, with or without doxycycline induction. (**E**) Representative immunofluorescence images showing subcellular localization and morphology of cells expressing NKX2-5 or the indicated deletion mutants. NKX2-5 signal is shown in green; merged images include DAPI and F-actin staining. (**F**) Pairwise correlation of transcriptional signatures induced by full-length NKX2-5 and indicated mutants. Summary tracks indicate relative protection, toxicity and localization patterns (N, nuclear; C, cytoplasmic) *in vitro*. (**G**) Transcriptional profiles of NKX2-5 variants and oxidative challenge. Heatmaps showing RNA-seq log_2_ fold changes for selected pathway genes after 4 days or 10 days of *in vitro* expression of GFP, full-length NKX2-5 (WT), or indicated NKX2-5 mutants relative to GFP control. The lower heatmap shows gene-expression changes after oxidative challenge relative to untreated controls. Genes are grouped by pathway annotation. Red indicates increased expression and blue indicates decreased expression. (**H**) GSEA pathway concordance between *in vitro* and *in vivo* NKX2-5ΔHD expression. Scatter plot comparing pathway-level enrichment signatures from RNA-seq analysis of NKX2-5ΔHD versus GFP *in vitro* and *in vivo*. Each point represents one GSEA pathway. Colored and labeled pathways indicate FDR < 0.05 in both comparisons. Pathways with FDR = 0 are shown at |score| = 5. Human gene sets were matched to mouse symbols without ortholog conversion. (**I**) *In vivo* pathway analysis of Old RPE vs Young RPE (n=4 each); NKX2-5ΔHD vs GFP (n=4 each); and full-length NKX2-5 vs GFP (10 weeks, n=3 each; 3 weeks, n=4 each) in aged RPE. Heatmap shows Hallmark preranked GSEA results from RPE RNA-seq after AAV-mediated expression of the indicated gene. Colors indicate normalized enrichment scores, with red indicating enrichment and blue indicating suppression. * FDR < 0.05 ** FDR < 0.01 *** FDR < 0.001 Data are shown as mean ± s.e.m.

To define the transcriptional programs associated with NKX2-5ΔHD-mediated oxidative resilience, we performed RNA-seq after 4 days of mutant expression and compared global transcriptional profiles (Fig. 5F) and curated path-way-associated genes (Fig. 5G). NKX2-5 and NKX2-5ΔHD induced a distinct transcriptional state marked by increased MHC-I and antioxidant gene expression, together with reduced hypoxia, TNFα/NF-κB, ECM, p53 apoptosis, and mTORC1/UPR-associated stress programs (Fig. 5G). These changes largely opposed the transcriptional responses induced by NaIO_3_-mediated oxidative stress.

Among the protective mutants, NKX2-5ΔHD uniquely preserved RPE identity genes while avoiding induction of matrix remodeling genes (*e*.*g*., *FN1* and *CAV1*) and inflammatory activation programs (*e*.*g*., *IL6* and *C3*). When expression was extended to 10 days, these protective transcriptional features were largely maintained by NKX2-5ΔHD, whereas full-length NKX2-5 showed stronger activation of stress-associated pathways (Fig. 5G).

The two YRD-disrupted mutants partially lost MHC-I activation and suppression of NF-κB/p53-apoptosis programs, while largely retaining suppression of hypoxia and mTORC1/UPR pathways, providing a potential molecular correlate of their partial loss of protection. In contrast, the N1-137 truncation abolished these transcriptional changes and resembled the GFP control, consistent with its loss of protective activity (Fig. 5G). Subretinal delivery of AAV2-CAG-NKX2-5_N1-137 to aged mice also failed to improve visual acuity (fig. S5C).

In addition to their effects on transcription, non-HD domains such as the YRD have been reported to mediate protein–protein interactions^68^. We therefore performed co-immunoprecipitation–mass spectrometry analyses of ΔTN, ΔYRD and YRD^Y-A^ mutants, and found reduced association of CSNK2A1 and CSNK2B with the ΔYRD and YRD^Y-A^ mutants, but not with the ΔTN (fig. S5, D to F). CSNK2A1 and CSNK2B genes encode casein kinase II (CK2) subunits, an essential enzyme involved in cell survival and proliferation^69,70^, and have been recently reported to control mitochondrial quality in cardiomyocytes^71^. Thus, protein interactions mediated by YRD may also partially contribute to oxidative resilience.

Although the N-terminal fragment lacked protective activity *in vitro* and *in vivo*, mRNA and protein measurements of the truncation mutants suggested that the N terminus influences *NKX2-5* transcription levels. For example, expression of the C-terminal fragment C198-324 (Δ1-197), or deletion of the AlphaFold-predicted N-terminal region spanning residues 14–41 (Δ14-41), resulted in low mRNA and protein levels. Conversely, the N-terminal fragment (N1-137) still maintained mRNA and protein levels (fig. S5, A and B).

These findings support a hypothetical working model in which distinct NKX2-5 regions mediate separable activities (fig. S5G). The N terminus contains determinants that promote *NKX2-5* mRNA and protein production or stability, whereas the homeodomain is linked to CARD10 association and the inflammatory, identity-loss phenotype that emerges with prolonged expression. In contrast, the YRD-containing C-terminal region supports oxidative resilience and is associated with CSNK2A1/CSNK2B interactions and protective transcriptional programs. This hypothetical model explains how homeodomain deletion may uncouple the beneficial resilience activity of NKX2-5 from the toxicity associated with ectopic overexpression.

### NKX2-5ΔHD partially reverses age-associated transcriptional programs *in vivo*

We next asked whether the protective transcriptional state induced by NKX2-5ΔHD *in vitro* was preserved *in vivo*. RNA-seq of aged RPE after AAV-mediated NKX2-5ΔHD expression revealed coordinated suppression of hypoxia, inflammatory, stress-response, EMT, glycolysis, and apoptosis-related pathways (Fig. 5H), consistent with the functional programs identified *in vitro*. These changes partially reversed age-associated pathway activation (Fig. 5I), and were distinct from prolonged full-length NKX2-5 expression, which showed greater induction of stress-associated programs.

Notably, a shorter, 3-week period of full-length NKX2-5 expression more closely resembled NKX2-5ΔHD (Fig. 5I), suggesting that homeodomain-associated stress responses emerge with prolonged expression. This temporal pattern is consistent with the visual acuity data, in which NKX2-5 initially improved function before the benefit began to diminish with longer treatment (Fig. 3I). Together, these data associate NKX2-5ΔHD-mediated functional improvement *in vivo* with suppression of several stress- and age-associated transcriptional programs that also reduced *in vitro*.

### Late-life systemic delivery of NKX2-5ΔHD improves grip strength and reduces frailty in male mice

To test whether NKX2-5ΔHD could promote systemic benefits beyond the retina, we delivered AAV9-CAG-d2EGFP or AAV9-CAG-NKX2-5ΔHD systemically to 25-month-old male C57BL/6 mice by retro-orbital injection and followed physiological function and frailty longitudinally; lifespan monitoring is ongoing (Fig. 6A). The use of only male mice reflects the limited availability of female mice at this advanced age and is consistent with prior AAV studies involving systemic delivery in aged mice ^29,30^.

**Fig. 6.**
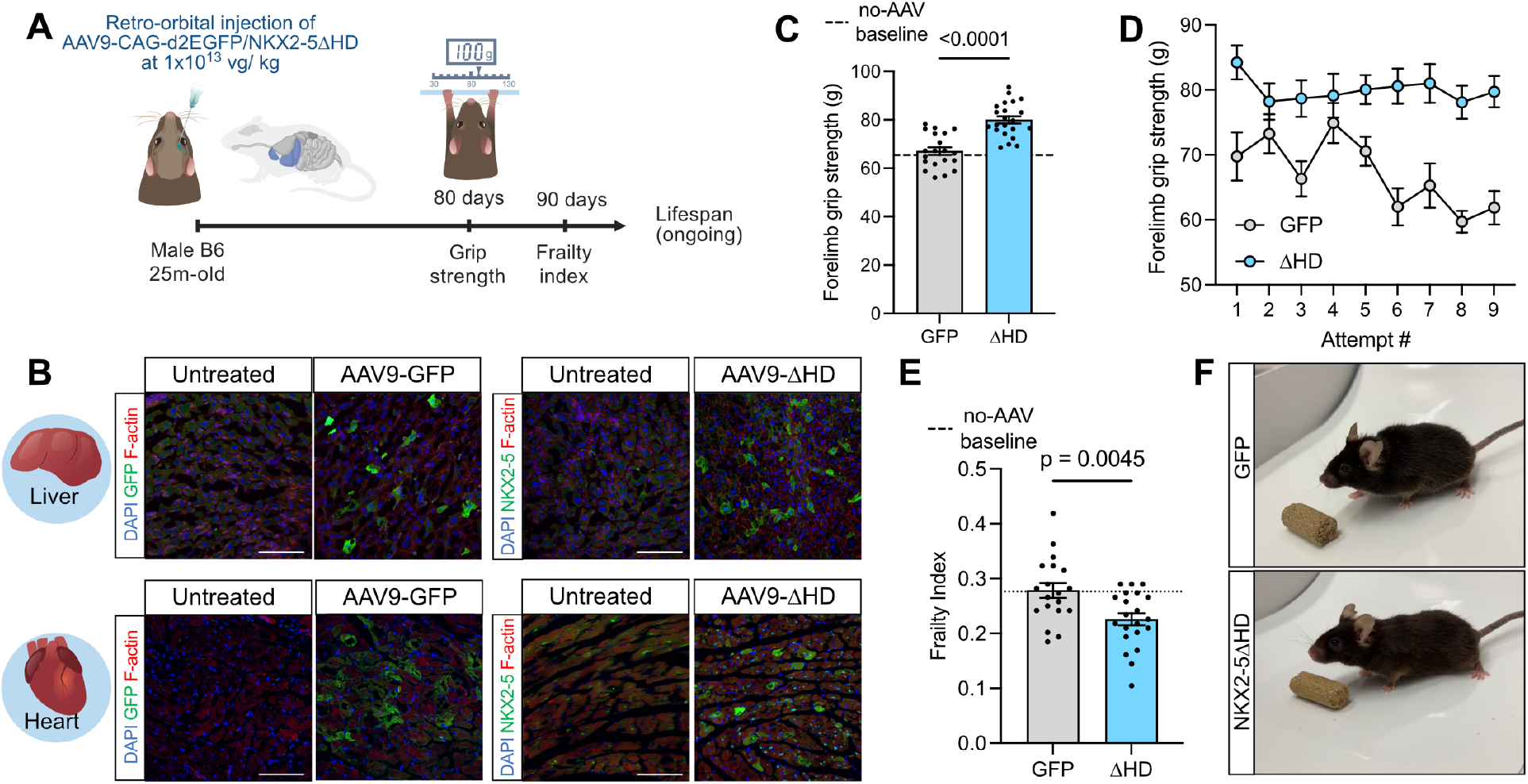
Systemic delivery of NKX2-5ΔHD improves grip strength and reduces frailty in aged male mice. (**A**) Systemic AAV-NKX2-5ΔHD treatment and longitudinal healthspan assessment in aged mice. Twenty-five-month-old male C57BL/6 mice received retro-orbital injection of AAV9-CAG-d2EGFP or AAV9-CAG-NKX2-5ΔHD at 1 × 10^13^ viral genomes per kg. Forelimb grip strength was assessed 80 days after injection, followed by frailty index scoring at 90 days. Mice are being monitored for lifespan, and the study is ongoing. (**B**) Representative immunofluorescence images of liver and heart tissues from untreated, AAV-GFP-treated, or AAV-NKX2-5ΔHD-treated aged male mice, stained for GFP or NKX2-5ΔHD expression, together with nuclear and tissue counterstains. Scale bar, 100μm. (**C, D**) Forelimb grip strength performance of AAV9-GFP- and AAV9-hNKX2-5ΔHD-treated male mice at 27.5 months of age, 80 days after AAV injection (GFP, n=20; NKX2-5ΔHD, n = 22). The dash line indicates forelimb grip strength of untreated age-matched controls (n = 4), with reference data shown in **Fig. S6C**. Panel C shows the mean grip strength across those attempts for each mouse; panel D shows performance across nine repeated attempts. Examiners were masked to group allocation. (**E**) Frailty index of AAV9-GFP- and AAV9-hNKX2-5ΔHD-treated male mice at 28 months of age, 90 days after AAV injection (GFP, n = 19; NKX2-5ΔHD, n = 21), one animal in each group died between 80 and 90 days after AAV injection. The dashed line indicates the mean of untreated age-matched controls (n = 4). Examiners were masked to group allocation. (**F**) Representative images of AAV9-GFP- and AAV9-hNKX2-5ΔHD-treated male mice selected based on frailty index values near their respective group means (0.266 and 0.202). The two group differences in kyphosis, grip strength, menace reflex, and grimace scale. Data are shown as mean ± s.e.m.

Immunostaining and transcriptomic analysis confirmed AAV9-mediated GFP or NKX2-5ΔHD expression in the liver and heart, two major organs efficiently transduced following systemic delivery (Fig. 6B and fig. S6, A and B), consistent with prior reports^29,72^. We then performed non-invasive healthspan assessments of these mice in a masked manner. At 80 days post-injection, NKX2-5ΔHD-treated mice not only showed significantly higher forelimb grip strength than d2EGFP-treated controls, measured by both average and peak grip strength of nine repeated attempts (Fig. 6C and fig. S6, C to E), but also maintained their performance across attempts (Fig. 6D). Although the mechanism underlying the improved grip strength remains unclear, particularly given the limited transduction of skeletal muscle by AAV9 at the given dose^21,72,73^ (fig. S6A), improved cardiac function and blood supply could potentially reduce exercise induced lactate accumulation, but this possibility warrants further investigation in future studies.

At 90 days post-injection, frailty was scored using a 31-item clinical index, and NKX2-5ΔHD-treated mice showed a reduced frailty index compared with GFP controls (Fig. 6E), similar to the effect reported with cyclic AAV9-mediated OSK expression^29^. Beyond grip strength, NKX2-5ΔHD also improved the menace reflex and showed trends toward reduced kyphosis and lower grimace scores, the latter reflecting facial expressions associated with pain or distress (Fig. 6F and fig. S6F). During the first 18 weeks, 126 days of treatment, continuous AAV9-NKX2-5ΔHD expression did not cause more body weight loss than GFP controls (fig. S6G). Although the lifespan experiment is still ongoing and its long-term safety and efficacy remain to be determined, the assessments so far indicate that systemic NKX2-5ΔHD delivery improves selected functional and frailty measures in aged male mice and warrant further investigation of benefits beyond the retina.

## Discussion

Aging is characterized not only by progressive functional decline but also by a loss of resilience to environmental and metabolic stressors. This decline is particularly relevant in the retinal pigmented epithelium (RPE), a long-lived and metabolically active tissue exposed to lifelong oxidative burden. Here, we established a genome-wide overexpression screening platform to systematically identify genes that enhance oxidative resilience in RPE cells. Unlike loss-of-function approaches, overexpression screening can interrogate the protective potential of the broader human ORFome regardless of whether a gene is normally expressed in RPE.

Our screening framework incorporated several methodological advantages for studying cellular resilience. We developed a replating-based selection strategy after observing that severely oxidatively damaged cells can remain attached while losing the capacity to reattach and proliferate. By adapting the logic of Death-seq^74^ to an overexpression screen, this approach enriches for genes that preserve functional viability after oxidative injury while improving signal-to-noise. The same framework could be extended to other cell types and disease-relevant stressors.

Among the protective genes identified, NKX2-5 revealed an unexpected capacity to promote oxidative resilience and improve age-associated phenotypes. NKX2-5 is best known as a cardiac developmental transcription factor required for heart formation and implicated in congenital heart disease, while its functions in adult tissues remain less understood^75^. Its activity in aged RPE suggests that developmental regulators can harbor stress-resilience functions outside their native lineage. This possibility is particularly intriguing because the heart and retina contain long-lived, metabolically active cells that must sustain function for decades under substantial oxidative demand. Cardiomyocytes sustain lifelong function under high metabolic and oxidative demand, pumping large volumes of oxygenated blood each day. This exceptional workload suggests that cardiomyocytes may possess intrinsic programs that support long-term oxidative resilience and cellular longevity.

Our domain-dissection studies further indicate that the protective activity of NKX2-5 can be separated from its canonical homeodomain-mediated function. Deletion of the homeodomain preserved oxidative protection while reducing the toxicity associated with full-length NKX2-5, supporting further evaluation of NKX2-5ΔHD as a therapeutic candidate. More broadly, these findings illustrate how multidomain transcription factors may contain distinct functional modules that can be dissected and engineered to retain beneficial stress-resilience activities while minimizing unwanted developmental or transcriptional effects.

A major challenge for reprogramming-based strategies, including transient OSKM expression and cyclic OSK treatment, is balancing efficacy with the potential toxicity of prolonged factor expression. We therefore extensively evaluated long-term NKX2-5ΔHD expression and found that this engineered single factor retained functional benefits with limited detectable toxicity, including after prolonged expression. This property may simplify therapeutic implementation.

The screen also revealed that not all forms of oxidative resilience are mechanistically equivalent. The LTBR/TNFα findings suggest that oxidative resilience in RPE can arise through inflammatory stress-response programs that carry important tradeoffs. Although TNF-family signaling is often viewed as pathogenic in AMD because it promotes NF-κB activation and EMT-like loss of RPE identity, our data suggest that this pathway may also function as an endogenous response mechanism to activate *SOD2* under oxidative stress (Fig. 2 and Fig. S2). These findings imply a plausible hypothesis that EMT-like remodeling may serve as a natural “escape” response, one that encourages cells to migrate out of harsh environments and activate endogenous protection, at the cost of loss of their original identity and function. This framework may help explain why indiscriminate suppression of TNFα signaling can, in some contexts, weaken intrinsic cytoprotective responses and exacerbate RPE death^76^. Combining pathway-specific interventions with antioxidant protection may therefore warrant consideration. In contrast, NKX2-5ΔHD provides a non-EMT route to oxidative resilience, highlighting the importance of identifying mechanisms that preserve both cell survival and cell identity.

The systemic effects of NKX2-5ΔHD further suggest that its protective activity may extend beyond the retina. In aged male mice, retro-orbital injection of AAV9-hNKX2-5ΔHD improved forelimb grip strength and reduced frailty. In our prior work, retro-orbital injection of AAV9-rtTA; TRE-OSK and cyclic expression of OSK also reduced frailty at late age, and led to lifespan extension^29^. These observations raise the possibility that the protective activity of NKX2-5 identified in RPE, may parallel OSK, reflects a broader stress-resilience program rather than a retina-specific effect.

Several considerations define the scope of our findings. First, ORF screens may miss factors requiring specific cofactors or cellular contexts, and our ARPE-19/NaIO_3_ screen represents one acute oxidative insult rather than the complex pathology of AMD. Second, although ectopic expression of factors such as NKX2-5 revealed unexpected protective activities, these experiments do not establish an endogenous role for these factors in the tissues where they are normally highly expressed. Likewise, transcriptomic age provides a molecular correlate of aging but, on its own, does not establish functional age reversal; therefore it needs to be interpreted together with functional and physiological measurements. Finally, our systemic NKX2-5ΔHD experiments were performed in late-aged male mice, and whether the effects generalize across sexes remains to be determined. The tissues and mechanisms underlying the improvements in grip strength and frailty also remain unresolved, as does whether these health benefits ultimately translate into increased lifespan.

NKX2-5 belongs to the evolutionarily conserved tinman/Nkx cardiac transcription factor family^77^, part of a regulatory network that predates vertebrates and accompanied the emergence of heart-like contractile organs more than 500 million years ago^78^. Although its canonical role in cardiogenesis depends on homeodomain-mediated transcription, ectopic expression in RPE uncovered a separable, non-canonical activity: deletion of the homeodomain reduced toxicity while preserving oxidative resilience and age-associated transcriptional benefits (fig. S6H). NKX2-5ΔHD could potentially be developed for AAV-based gene therapy, used to engineer more resilient iPSC-derived RPE for cell replacement, or exploited to identify downstream druggable pathways. More broadly, this study suggests that ancient developmental regulators may harbor latent protective activities outside their native lineage, and genome-scale gain-of-function screening may provide a systematic route to discovering and engineering resilience programs to counter age-related degeneration in the retina and beyond.

## Supporting information

Supplementary Discussion

## Declaration of use of generative AI

The authors used ChatGPT for manuscript proofreading, then reviewed and edited the content as needed and took full responsibility for it.

## Acknowledgements

We are grateful to Caitlin Rausch for assistance with illustrations. We thank Stephen Elledge and Mamie Li for sharing the ORF library. We thank Tessa Bertozzi, Yue He, Julia Joung, Pu Zheng, Anna Puszynska, Mohamed EIBrolosy, Edwin Neumann, George Bell and all members of the Weissman lab for helpful assistance and discussions. We thank Constance Cepko, Patricia D’Amore, Jason Buenrostro, James Handa, Margarete Karg, Daisy Shu, Andy Zhao, Miele Macmillan, Caitlin Keiper and Yoshiko Iwamoto for their technical support and advice, and Yechen Shen, Chang Luo, and Ming Lei for fellowship support. We thank the Whitehead sequencing, bioinformatics, and imaging cores for their support. Y.R.L. was supported by postdoctoral fellowships from Glenn/AFAR and Life Sciences Research Foundation, and the NIH Pathway to Independence Award (K99EY037340). H.S. received support from the First Affiliated Hospital with Nanjing Medical University. S.S. was supported by Bright Focus AMD Fellowship. We thank NIH R01EY031748 and R01EY037682 (C.B.R), NIH P30EY005722 (to Duke), FFB Free Family AMD Award (to C.B.R.); Research to Prevent Blindness grant (to Duke), P30EY003790 (to Schepens Morphology Core), R01EY036910 (to B.R.K. and M.G.K.), R01EY025794, P01AG071463 (to B.R.K). V.N.G. was supported by NIA and Hevolution. J.S.W. is an H.H.M.I. investigator. We especially thank the Milky Way Research Foundation for the generous support (to J.S.W.).

## Author contributions

Y.R.L., and J.S.W. conceived the study. Y.R.L designed all experiments.

Y.R.L and K.K. performed the ORF screen, and K.K. led the analysis. Y.R.L. performed candidate validation *in vitro* and *in vivo*, with assistance from J.C.C., J.A. and H.S., under supervision of B.R.K. and J.S.W.. J.C.C., J.A., X.L., conducted cell culture and molecular cloning. Y.H. and Y.H.C assisted with transcriptomic analysis. Y.H.C., L.G., and M.M. support co-IP-MS experiment and analysis. S.H. and Y.H.C. performed AlphaFold analysis on NKX2-5 mutants. H.S. led the subretinal injection and OCT analysis. S.S., Z.C., Q.L., T.D., M.B., J.C. conducted mouse experiments and analysis. J.R., R.S., K.B. shared iPSC-RPE cells and reagents. A.T., and V.N.G. assisted with transcriptomic age analysis. M.G.-K., M.W.C., C.B.R. provided assistance to mouse experimental design. Y.R.L, B.R.K and J.S.W. co-wrote the manuscript with inputs from co-authors.

## Materials and Methods

### Data and materials availability

The ORF screen and RNA sequencing data will be deposited to GEO. Code will be available through GitHub. The mass spectrometry proteomics data have been deposited to the ProteomeXchange Consortium via the PRIDE 80 partner repository with the dataset identifier PXD084197.

Other data and material generated are available from the corresponding authors upon request. All figures were screened with the Proofig platform as part of an automated image-integrity assessment.

### Library amplification and annotation

The pooled ORF library used in this study was Library 2 published in Sack et al. ^17^, which consisted of a subset of ORFs cloned into lentiviral backbones (Ultimate ORF 24-mer ORFs and cherry-picked subset of ORFeome v8.1 30-mer ORFs). A similar subset library (using only the 24-mer library) was resequenced to confirm barcodes to ORFs in Legut et al. ^18^. We generated a consolidated ORF reference table for all read assignment and enrichment analyses. We used the newer confirmed reference from Legut et al and appended the cherry-picked subset. Beginning from a curated reference of 12,999 unique ORFs, we identified 736 barcodes that were assigned to more than one ORF in the source annotations ^18^; these ambiguous barcodes were removed. ORF names and symbols were then updated against the HGNC gene set; 184 ORFs absent from the HGNC set were resolved manually and assigned updated gene symbols. ORF lengths were taken from the Sack et al. reference where available; for ORFs lacking an annotated length, lengths were retrieved by querying UniProt. For the remaining subset without associated UniProt identifiers, lengths were annotated manually where they could be determined and otherwise recorded as missing (NaN). The final consolidated reference table can be requested from corresponding authors.

### Lentivirus production and functional titering

Lentiviral particles were produced in HEK293T C17 cells using third-generation packaging. HEK293T C17 cells were seeded 24 h prior to transfection in gelatin-coated T175 or T75 flasks with antibiotic free growth medium to improve transfection efficiency, and were transfected at ~70–85% confluency. Transfer plasmids included a pooled ORF library cloned into a pHAGE lentiviral backbone. Packaging plasmids encoding Gag/Pol, Rev, and VSV-G were co-transfected using FuGENE (Promega) in Opti-MEM according to the manufacturer’s instructions, maintaining a 3:1 FuGENE-to-DNA mass ratio. Fourteen to eighteen hours post-transfection, the medium was replaced with ARPE growth medium with antibiotics. Viral supernatants were collected 48–72 h post-transfection and clarified by low-speed centrifugation. Virus was concentrated by polyethylene glycol–based precipitation (Lenti-X Concentrator) at 4 °C followed by centrifugation at 1,500 × g for 45 min. Viral pellets were resuspended in ARPE growth medium and aliquoted. Concentrated lentivirus was stored at −80 °C until use. Functional viral titers were determined empirically using a GFP reporter virus. ARPE-19 cells were seeded at 5 × 10^4^ cells per well in 24-well plates and transduced the following day with increasing volumes of GFP-expressing lentivirus in the presence of 8 μg mL^−1^ polybrene. Twenty-four hours posttransduction, GFP expression was assessed by flow cytometry, and the percentage of GFP-positive cells was quantified. Based on these measurements, viral input volumes yielding ~30–40% transduction efficiency were identified and used to guide downstream infections, with the goal of minimizing multiple lentiviral integrations per cell.

### Pooled ORF library screening

For pooled ORF library screening, ARPE-19 cells, at passage 25, were expanded to 160 million cells and seeded into T175 gelatin-coated flasks. After 24 hrs of settlement, a pooled library of lentivirus was added to the cells. The amount of lentivirus used for transduction was titrated to result in 35% transduction efficiency to minimize potential of multiple ORF incorporation into a given cell. Plates were gently agitated to ensure even distribution of virus. 24 hours post-transduction, medium was replaced with fresh ARPE growth medium. 48 hours post-transduction, the cells were trypsinized and replated into new gelatin pre-coated T175 flasks with growth medium containing puromycin selection (8 μg mL^−1^), the untransduced cells would die and float and medium were replaced with new growth medium containing puromycin after 48 hrs of selection. The total puromycin selection was 3 days to enrich for transduced cells and allow cells to grow into confluency.

At this point, two replicates of cells maintaining 4000x coverage were collected after the cells were puromycin selected while subsequent cells were reseeded into experimental and control screen arms, two replicates each.

The experimental arm received sodium iodate (5mM NaIO_3_) treatment in serum-free ARPE medium for 18 hours. Afterwards, the detached (non-NaIO_3_ resistant dead cells) were collected, while the resistant live cells were reseeded to fresh ARPE medium containing serum and a population of cells maintaining 1000x coverage was collected. Matching the timeline, cells in the control arm were switched to serum-free starvation medium for 18 hrs and reseeded to serum-containing medium, and a population of cells maintaining 1000x coverage was collected.

Genomic DNA was isolated from samples collected at all time points using the NucleoSpin Blood L kit (Macherey-Nagel, 740954.20). ORF barcodes were amplified by two rounds of PCR using KAPA HiFi HotStart ReadyMix (Roche, 07958927001), followed by a third PCR (PCR3) to add Illumina sequencing adapters as previously described^17^. PCR3 products were gel-purified using the NucleoSpin Gel and PCR Clean-up kit (Macherey-Nagel, 740609.250). Libraries were sequenced on an Illumina NovaSeq 6000 using a 30-bp read with the heyME19 sequencing primer. Sequencing data is available through GEO.

### Pooled ORF screen analysis

Raw FASTQ files were pre-processed to extract barcode-containing reads. For each sequencing file, reads corresponding to the barcode region were isolated by selecting the appropriate read lines and truncating to a fixed barcode length. Identical preprocessing was applied across all samples, and trimmed barcode reads were saved as index-specific files. Because each biological sample was sequenced across two lanes, lane-level files (denoted “A” and “B”) were merged prior to downstream analysis, resulting in a single consolidated file per index. Index assignments corresponded to plasmid library, transduced populations, baseline populations, and live or dead cell fractions collected 18 h after NaIO_3_ treatment, each with two biological replicates. Trimmed barcode reads were mapped to the reference ORF–barcode library to assign barcodes to their corresponding ORFs. Mapping and downstream processing were performed using custom analysis scripts implemented in Python and R. For each sample, barcode counts were aggregated, and lane-collapsed counts were used for all subsequent analyses. Enrichment analyses were performed at both the barcode and gene (ORF) level using MAGeCK. Barcode- and gene-level count tables were normalized by total read depth. For gene-level analyses, barcode counts were first collapsed by ORF prior to normalization. Paired analyses were used where applicable to account for matched biological replicates. Differential enrichment was assessed across multiple biologically relevant comparisons, including live versus dead cells following NaIO_3_ treatment, NaIO_3_-treated versus baseline populations, baseline versus transduced populations, and transduced versus plasmid library controls. For each comparison, MAGeCK was run using paired replicates when appropriate, with gene-level log_2_ fold changes calculated using the mean fold-change method. ORFs were classified as enriched or depleted based on MAGeCK statistical outputs. Specific commands and analyses will be available on Github.

### Animals

All animal procedures were reviewed and approved by the Institutional Animal Care and Use Committees (IACUCs) at the Schepens Eye Research Institute of Massachusetts Eye and Ear and were conducted in accordance with institutional animal welfare guidelines. C57BL/6 mice were used for all studies, and obtained from the NIA aged rodent colonies in equal sexes for vision experiments. For the systemic effect experiment, 24-month-old male mice were obtained from the National Institute on Aging, while females at this age were not available. Mice were acclimatized for four weeks after arrival at the facility and before baseline measurements and retro-orbital injections. Mice were maintained under a 12-h light/dark cycle (6:00/18:00), at an ambient temperature of 21–22 °C and 40–50% humidity.

### Animal Experimental Design

Sample sizes were determined based on the standard deviations observed in our previous studies ^44^ using a two-sided t-test power analysis in G*Power (α = 0.05, power = 80%), with an additional 20% allowance for potential injection failure. Except for the retroorbital injection experiments in Fig. 6 where the use of only male mice reflects the limited availability of female mice at this advanced age, equal numbers of male and female mice were allocated before injection to account for possible sex-dependent effects. Mice exhibiting intravitreal bleeding or signs of inflammation, including corneal clouding or edema, during or after subretinal injection were excluded according to predefined criteria. When all injections were successful, additional animals were retained for functional analyses. Investigators performing the experiments and outcome assessments were masked to group allocation. Details regarding the ages and numbers of mice used in each experiment are provided in the corresponding figures and figure legends.

### Lentiviral and AAV vector cloning

Doxycycline-inducible lentiviral constructs were generated by Gateway LR recombination between pLIX_403 (Addgene #41395) and pDonor-ORF vectors, obtained either from the hORF Lite Collection (A11613, Thermo Fisher Scientific) or by synthesis through IDT/Azenta. The AAV constructs pAAV-CAG-d2EGFP, pAAV-CAG-hNKX2-5, and pAAV-CAG-hNKX2-5ΔHD were generated by Gateway LR recombination between in-house pDonor vectors and the pAAV-Gateway backbone (Addgene #32671).

### Mice Anesthesia

Mice were anesthetized with intraperitoneal ketamine/xylazine (100–200 mg/kg and 20 mg/kg, respectively) and topical 0.5% proparacaine (Bausch & Lomb). To prevent corneal dryness, GenTeal (Alcon) was applied after anesthesia. Following the procedure, aged mice received intraperitoneal Atipamezole hydrochloride (1.0 mg/kg) to reverse xylazine anesthesia and promote recovery.

### Subretinal delivery of AAV

15-month-old mice were anesthetized and subjected to subretinal AAV injection as previously described^44^. Briefly, pupils were dilated with one drop each of 1% tropicamide and 2.5% phenylephrine ophthalmic solution (PINE Pharmaceuticals, New York, USA). After adequate dilation, a sclerotomy was created approximately 1 mm posterior to the limbus in the superior-temporal quadrant using a 30-gauge needle. GONAK 2.5% hypromellose ophthalmic demulcent solution (AKORN, Illinois, USA) was applied to the corneal surface, and a 5-mm diameter contact lens (Warner Instruments, Connecticut, USA) was positioned to visualize the fundus under a surgical microscope.

A 36-gauge blunt needle (World Precision Instruments, Florida, USA) connected to a 10-μL NanoFil microsyringe injection system (World Precision Instruments, Florida, USA) was inserted through the preformed entry site and advanced into the subretinal space in the nasal-ventral mid-peripheral retina. A total of 1 μL AAV2 vector (1 × 10^9^ vg/eye) was injected into the subretinal space. Successful delivery was confirmed by the formation of a visible subretinal bleb. Following injection, excess hypromellose solution was gently removed with an eye spear. To minimize postoperative infection, neomycin/polymyxin B/bacitracin zinc ophthalmic ointment (Bausch + Lomb, Quebec, Canada) was applied to the injection site.

### OptoDrum Optomotor Response (OMR) Assessment

Visual acuity was assessed using an automated OptoDrum optomotor reflex (OMR) system^81^. Briefly, mice were placed on a central platform surrounded by four monitors displaying rotating vertical black-and-white sinusoidal gratings. The OptoDrum system used automated software-based tracking of the mouse outline, nose, and tail, and scored responses only when the animal was stationary, thereby reducing observer bias and increasing assay stringency. In addition, the system dynamically adjusted grating spacing according to head position, enabling more accurate spatial frequency presentation than traditional Optomotor.

During testing, contrast was maintained at 99.27% and rotation speed at 12°/s. Spatial frequency thresholds were determined using a predefined staircase paradigm ranging from 0.222 to 0.416 cycles/degree. Visual acuity results are reported as cycles per degree. Confirmation of the final threshold required two positive responses followed by three negative responses at the next higher spatial frequency. OptoDrum assay is more stringent than the classic Optomotor assay because it uses automated camera tracking, and dynamically adjusts stimulus spacing based on head position.

### Optical Coherence Tomography (OCT)

Optical coherence tomography (OCT) imaging was performed using a Bioptigen Envisu R-Class OCT system (Leica Microsystems). Mice were anesthetized and pupils were dilated before examination. Volumetric retinal scans were acquired using Bioptigen InVivoVue™ 2.4 software (1.4 × 1.4 mm scan area, 1.79 mm depth, 1000 A-scans/B-scan, 100 B-scans/volume, and 3-frame averaging). Images were analyzed with InVivoVue Diver 3.4.4 software, and retinal thickness was quantified automatically from the inner limiting membrane (ILM) to the retinal pigment epithelium (RPE).

### Electroretinogram (ERG)

ERG was conducted to evaluate the retinal function as previously described ^82^. In brief, mice were dark-adapted overnight and maintained under dim red light throughout the procedure on a built-in warming platform to preserve body temperature at 37 °C. Both eyes were dilated with 1% tropicamide and lubricated with Genteal (Novartis Pharma AG, Basel, Switzerland) before the stimulators were placed on the ocular surface. ERG a- and b-wave responses were recorded at flash intensities of 0.01, 0.1, and 1 cd·s/m^2^, whereas c-wave responses were measured at 24.1 cd·s/m^2^. Data were analyzed using CELERIS software (Diagnosys). The a-wave amplitude and implicit time were measured from baseline to the trough of the initial negative deflection, the b-wave amplitude from the a-wave trough to the peak of the subsequent positive deflection, and the c-wave amplitude from baseline to the maximal positive peak.

### Multifocal Electroretinography (mfERG)

MfERG, a sensitive method for assessing localized retinal function, was performed as previously described ^83^ using the Celeris pattern stimulator and Diagnosys mfERG software v3.8.1 (Diagnosys LLC, Lowell, MA, USA). Briefly, mice were dark-adapted overnight before testing and kept under dim red light during the procedure. Following pupil dilation with 1% tropicamide for 15 min, Genteal lubricant (Novartis Pharma AG, Basel, Switzerland) was applied to both eyes to maintain corneal hydration and optical contact. The pattern stimulator examined a 50° retinal field containing 7 hexagons was positioned on the ocular surface of the experimental eye and a flash stimulator was placed on the contralateral eye. mfERG responses were analyzed for each hexagon (1–7) using a linear mixed-effects model with Animal ID as the subject identifier, Group as the between subject factor, and Hexagon as the within subject factor. Post hoc comparisons were performed using estimated marginal means with Holm correction for multiple testing. All tests were two-sided, and p < 0.05 was considered statistically significant.

### Establishment of Stable ARPE-19 Cell Lines with Doxycycline-Inducible Expression

Doxycycline-inducible constructs were generated by Gateway LR recombination of pDonor-ORF inserts into pLIX_403 (Addgene #41395). The pDonor clones were obtained from the Ultimate hORF (Lite) Collection (Thermo Fisher Scientific, A11613); the NKX2-5 mutant and deletion ORFs were synthesized by IDT and Azenta. To establish stable lines, ARPE-19 cells were plated on day 0, transduced with lentivirus and 8 μg/mL polybrene on day 1, and given fresh medium on day 2. Selection with 4 μg/mL puromycin began on day 4 and was typically complete by day 8, when all uninfected control cells had died. Selected cells were expanded and plated into wells from day 9. After reaching confluence, the cells were maintained in nicotinamide-containing maturation medium for three weeks, followed by induction with 2 μg/mL doxycycline.

### CRISPR Knockout

Dual guide RNAs targeting exons of genes of interest were picked from human Brunello library ^84^ (sequences listed below). They were cloned into lentiCRISPRv2-Opti by Golden Gate assembly using the NEBridge Golden Gate Assembly Kit (BsmBI-v2), following the manufacturer’s instructions. Constructed plasmids were transformed into NEB Stable Competent *E. coli*. Single clones were picked and cultured, and plasmids were isolated by miniprep for sequencing confirmation.

For lentivirus production, HEK293T cells were co-transfected using TransIT transfection reagent with the sequence-verified transfer vector (2 μg) and the packaging plasmids pMDL (1.3 μg), pRSV-Rev (0.5 μg), and pMD2.G (VSVG; 0.7 μg). The medium was replaced the next day with ViralBoost Reagent. Viral supernatants were harvested 48 h after transfection, centrifuged at 500 × g for 5 min to remove cell debris, and stored at −80 °C. Target cells were seeded at 2 × 10^5^ cells per well in six-well plates 48 h before transduction. Cells were incubated with 1 mL viral supernatant, 8 μg/mL polybrene, and 1 mL culture medium, and the medium was replaced with fresh medium the following day. Approximately 96 h after transduction, the cells were passaged into medium containing 4 μg/mL puromycin; selection was maintained for five days. Untransduced cells were treated with puromycin as a selection control, and their death confirmed the effectiveness of selection.

### Replate and Counting

After three weeks of maturation, ARPE-19 cells were induced with 2 μg/mL doxycycline in maturation medium for 3 days. Cells were then cultured for an additional day in starvation medium containing 2 μg/mL doxycycline before exposure to the indicated oxidative stress conditions in starvation medium. Two wells per group received no oxidative stress treatment and served as untreated controls for normalization of cell counts. Bright-field images were acquired using an Incucyte system.

For the LDH assay, 40 μL of culture supernatant was mixed with LDH assay reagent and incubated for 30 min at room temperature protected from light. Stop solution was then added, and absorbance was measured at 492 nm using a microplate reader. Baseline LDH release was defined using cells cultured without DOX induction or NaIO_3_ treatment, whereas maximum LDH release was determined using cells treated with lysis solution. LDH release was normalized between these baseline and maximum-damage controls and converted to membrane integrity. Accordingly, membrane integrity >100% indicates LDH release below baseline.

For the replating survival assay, cells were trypsinized, replated in growth medium, and cultured for 1 day. To minimize cell drying during replating, individual medium changes and wash steps were completed within approximately 7 s using gentle pipetting. Unattached cells and debris were removed by PBS washing before bright-field imaging. Cells were then harvested, resuspended in 110 μL PBS containing 5% FBS, and analyzed using an Attune Flow Cytometer, with 70 μL acquired per sample.

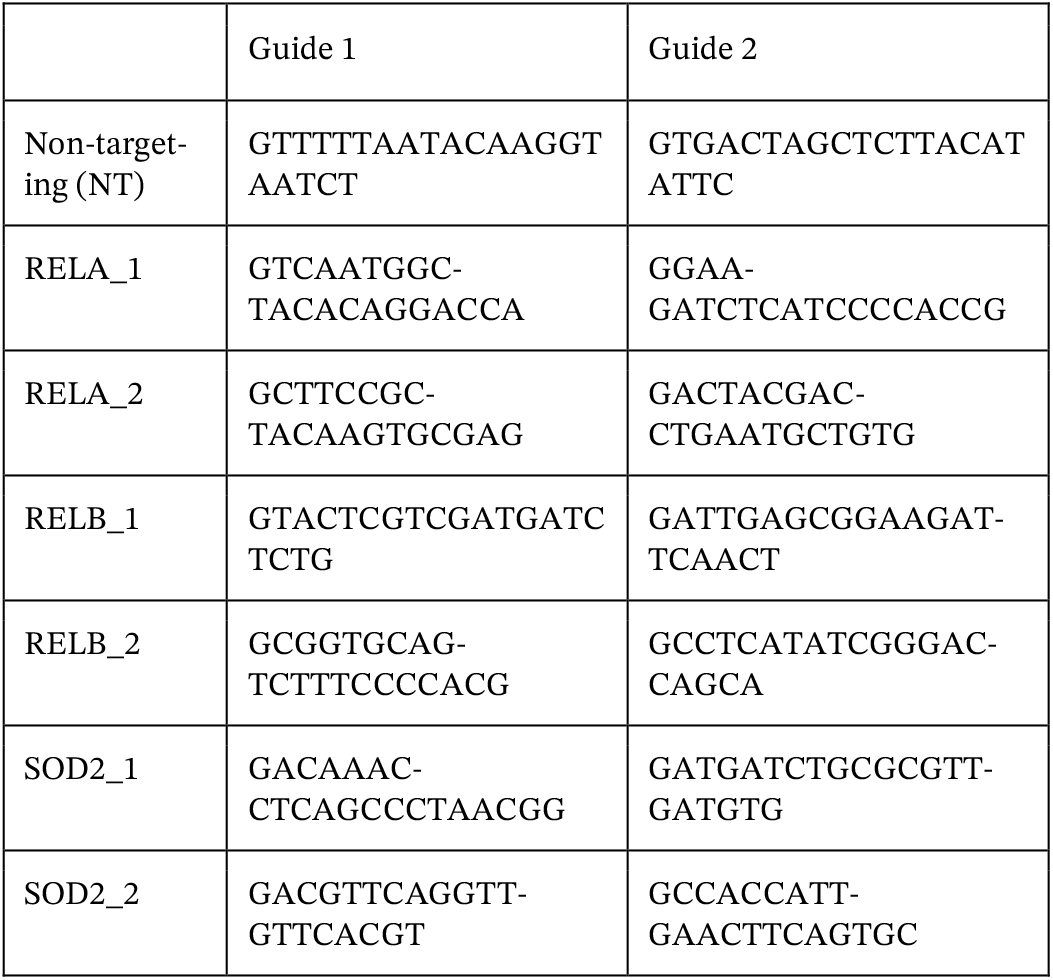

### Complement stress assay

Matured ARPE-19 cells were treated with 30% Complement Competent Human Serum (CC-HS) in maturation medium with or without 2 μg/mL doxycycline (MP Biomedicals, 198955) for 7 days and then the medium was collected and subjected to the desired assay. The maturation medium containing 30% CC-HS was refreshed at day 3 and day 5 during the treatment process. Damage was normalized to LDH release from lysed cells in the undamaged control. Because LDH release during CC-HS treatment accumulates over 48 hours, normalized values can exceed 100%.

### High-throughput affinity-enrichment mass spectrometry and analysis

For preparation of lysates, four biological replicates of stable ARPE-19 cell lines expressing the indicated ALFA-tagged proteins were generated by lentiviral transduction and selection with puromycin as described above. ALFA-tag affinity enrichment and proteomic sample preparation were performed as described previously ^85^. In brief, cells were trypsinized, pelleted, and lysed at 4 °C in IP lysis buffer (0.5% IGEPAL CA-630, 50 mM Tris-HCl pH 7.5150 mM NaCl, 5% glycerol) freshly supplemented with cOmplete EDTA-free protease inhibitor (Roche, 05056489001), 0.1% (v/v) in-house benzonase, and 1 mM MgCl_2_. Lysates were cleared by centrifugation at 4,000 × g for 15 min at 4 °C and flash frozen.

Affinity enrichment on clarified lysates was performed with streptavidin-coated 384-well plates coated with biotinylated anti-ALFA nanobody (NanoTag, N1505-Biotin, or produced in house). Cleared lysate was transferred to the nanobody-coated plate at 50 μL per well, diluted to 125 ng per well in 50 μL volume, incubated for 2 h at 25 °C and 1,200 r.p.m., and washed three times with ice-cold wash buffer (50 mM Tris-HCl pH 7.5150 mM NaCl, 5% glycerol). Peptides were acidified, desalted using Evotips, and analyzed by LC-MS/MS.

Raw dia-PASEF files were processed with DIA-NN 2.2.0. All 21 runs were analyzed together in a single search against an in silico spectral library predicted by deep learning with DIA-NN 2.2.0 from two sequence databases: a project-specific FASTA containing the ALFA-tagged NKX2-5 constructs and related bait sequences (20200603_IP-MS_seq.fa) and the human UniProt/SwissProt reference proteome (release 2024-07-22), together yielding 4,553,781 precursors covering 20,431 proteins and 20,215 genes. In silico digestion allowed cleavage C-terminal to lysine and arginine including before proline (--cut K*,R*) with up to one missed cleavage and N-terminal methionine excision, peptide length 7-30, precursor charge 1-4, precursor m/z 300-1800 and fragment m/z 200-1800. Carbamidomethylation of cysteine was a fixed modification; no variable modifications were used. The search was run with a scan window of 7, mass accuracies fixed at 12 ppm (MS2) and 15 ppm (MS1), peptidoform scoring, match-between-runs (--re-analyse) and retention-time profiling enabled. Precursor and protein identifications were filtered at 1% FDR and protein quantities were calculated with QuantUMS/MaxLFQ.

### Live cell staining

For ORF induction, matured ARPE-19 cells (three weeks in maturation medium) were initially cultured in maturation medium with 2 μg/mL doxycycline for 3 days. The cells were then switched to starvation medium containing 2 μg/mL doxycycline for an additional day prior to exposure to 5 mM NaIO_3_ in starvation medium for 4h. The cells were washed once with 1×DPBS and incubated at 37 °C for 30 minutes in the dark with freshly prepared staining buffer containing 1 μM H2DCFDA (Invitrogen, D399), and 200 nM MitoTracker Deep Red (Invitrogen, M46753). The cells were then washed again with 1×DPBS to remove residual staining buffer, harvested and resuspended in 110 uL 1×DPBS supplemented with 5% FBS and 1 μg/mL DAPI. Samples were analyzed using an Attune Flow Cytometer and a volume of 70uL per sample was collected.

### RPE and Retinal Flatmount and Immunofluorescence

Mouse eyes were enucleated and fixed in 4% paraformaldehyde (PFA) at 4 °C overnight. After removing the anterior segment, the neuroretina was gently peeled away from the underlying RPE–choroid cup, which was then radial-cut 7–8 times to generate an RPE or retinal flatmount. Dissected retinas and RPE flatmounts were washed in PBS before being blocked with BlockAid Blocking solution (Thermo, B10710) with 0.1% Triton X-100 for one hour. Then flatmounts were stained overnight with primary antibodies at 4 °C and then secondary antibodies at room temperature for 2 h. Between changes of solutions, all flatmounts were washed 3 times, for 5 min each time. Antibodies used were as follows: goat anti-NKX2-5 (R&D systems, AF2444), and Phalloidin-647 (for F-actin, Thermo, A22287). All retina and RPE flatmounts were mounted with Vectashield Antifade Mounting Medium.

### Western Blotting

Cells were lysed in ice-cold RIPA buffer supplemented with protease inhibitors. After clarification, lysate concentrations were determined by BCA assay. Samples containing 15–25 μg protein per lane were mixed with Laemmli buffer, boiled at 95 °C for 5 min, and separated by SDS-PAGE in MOPS running buffer at 150 V. Proteins were transferred to nitrocellulose membranes using the Trans-Blot Turbo semi-dry system. Following Ponceau S staining, membranes were blocked for 1 h at room temperature in EveryBlot blocking buffer, incubated with primary antibodies overnight at 4 °C, followed by secondary antibodies for 1 h at room temperature. Chemiluminescent signals were captured using an automated blot imaging system. Antibodies used were as follows: mouse anti-actin HRP (Abcam, ab49900); goat anti-NKX2-5 (R&D systems, AF2444); mouse anti-FLAG (MilliporeSigma, F1804); rabbit anti-vinculin (Cell Signaling Technology, 13901).

### Mouse RPE Isolation

Enucleated eyes were kept in cold PBS on ice. After the anterior segments were removed, the retinas were either collected separately and snap-frozen or discarded. The remaining posterior eyecups were briefly rinsed with cold PBS and then placed in 200 μL RNAprotect Cell Reagent (Qiagen) for 10– 30 min at room temperature. Tubes were agitated to release RPE cells, and the eyecups were removed. RPE cells were centrifuged at 2,500 rpm for 5 min at room temperature, resuspended in 200 μL TRIzol, and flash-frozen on dry ice. RPE purity was assessed by RNA-seq: samples showed high Rpe65 expression and low expression of Cx3cr1, Cdh5, and Rho, markers of microglia, endothelial cells, and photoreceptors, respectively.

### RNA-seq Library Preparation and Paired-End Sequencing

Total RNA was isolated with TRIzol Reagent (Thermo Fisher) supplemented with 1 μL glycogen (10 mg/mL) to improve yield. RNA quantity was measured using a Qubit 3.0 Fluorometer (Life Technologies), and integrity was assessed with an Agilent TapeStation. Genewiz prepared the RNA-seq libraries using the SMART-Seq v4 Ultra Low Input Kit (Clontech) for fulllength cDNA synthesis and Nextera XT (Illumina) for library construction. Following limited-cycle PCR indexing, library quality was confirmed by Qubit and TapeStation. Multiplexed libraries were clustered on two lanes of an Illumina HiSeq flow cell and sequenced as 2 × 150 bp paired-end reads. HiSeq Control Software was used for image analysis and base calling; bcl2fastq v2.17 converted BCL files to FASTQ and demultiplexed reads, permitting one mismatch in index identification.

### RNA-seq Analysis

Paired-end human and mouse reads were mapped to the canonical chromosomes of the GRCh38/hg38 or GRCm38/mm10 genomes, respectively, with STAR (v 2.7.1a)^86^ using a genome index created using Ensembl 96 gene models (with sjdbOverhang=100). Gene counts were obtained for all genes using featureCounts (v1.6.2)^87^ with the same GTF file as for STAR and options “-p -s 1”. Differential expression analysis was performed using the DESeq2 framework. For analyses conducted in R (v3.6.3), DESeq2 (v1.26.0)^88^ was used with independentFiltering=FALSE and normal log_2_-fold-change shrinkage. For analyses implemented in Python, raw integer count matrices were analyzed using PyDESeq2 ^89^ after dataset-specific filtering of low-count genes. Sample-specific size factors were estimated using the median-of-ratios method, and negative-binomial generalized linear models were fitted using the design specified for each experiment. Reported log_2_ fold changes represent fitted contrast coefficients from these models.

### Sample quality assessment and pathway expression scores

Principal component analysis (PCA) of log_2_-transformed, size-factor-normalized counts was used for exploratory sample quality assessment. In the NKX2-5 variant dataset, GFP-a was visualized as an outlier in the pre-exclusion PCA and formally flagged using a robust sample-correlation criterion based on each sample’s median Spearman correlation with all other samples. The primary analysis excluded GFP-a, and a sensitivity analysis including all four GFP samples confirmed that its inclusion did not materially alter the conclusions.

For pathway-by-sample expression scores, genes passing the dataset-specific expression filter were retained and normalized using DESeq2 size factors. Normalized expression was centered and scaled separately for each gene across samples using the sample standard deviation, without log transformation. For each pathway and sample, the score was calculated as the mean standardized expression of the available pathway-member genes. These scores were used as descriptive expression summaries rather than formal tests of pathway activity.

For correlation analyses relating gene expression to cell survival, gene counts were normalized using DESeq2-estimated size factors before calculating correlations.

### Transcriptomic signature analysis

To identify transcriptional programs associated with aging and NKX2-5 or NKX2-5ΔHD expression in RPE, differential expression analysis was additionally performed using edgeR for comparisons between (1) young and old control RPE and (2) old GFP-treated and old NKX2-5- or NKX2-5ΔHD-treated RPE. Genes were ranked using a signed log-transformed P value: −log(pv)×sgn(lfc), where *P* and logFC represent the P value and log fold change for each gene, respectively.

Gene set enrichment analysis (GSEA) was performed on the resulting preranked gene lists using the fgsea package in R with 10,000 permutations and multilevel Monte Carlo sampling. Gene sets were obtained from the HALLMARK, KEGG, and REACTOME collections in the Molecular Signatures Database (MSigDB). P values were adjusted for multiple testing using the Benjamini-Hochberg procedure, and adjusted *P* < 0.05 was considered significant.

### Transcriptomic clock analysis

Before applying transcriptomic clocks, human ARPE-19 or mouse RPE RNA-seq data underwent filtering and normalization. Genes with fewer than 10 reads in more than 80% of samples were excluded. The filtered data were then processed with Relative Log Expression (RLE) normalization, log-transformation, and YuGene transformation. Missing expression values for clock genes not detected in the dataset were imputed using their corresponding precomputed average values. Normalized gene expression profiles were centered to the median profile of old control samples. Transcriptomic age (tAge) for each sample was estimated using either rodent (for mouse RPE) or multi-species (for ARPE-19) Bayesian Ridge multi-tissue transcriptomic clocks of chronological age ^47^. Differences in mean tAge between young and old controls, as well as between old controls and age-matched NKX2-5-overexpressing samples, were evaluated using a mixed-effects ANOVA model implemented via the rma.uni function from the metafor package in R. Module-specific rodent and multi-species transcriptomic clocks of chronological age were applied to scaled relative gene expression profiles using the same framework. For each module, tAges were standardized across samples, and then differences in average standardized tAge between old control and treated groups were assessed with ANOVA. Resulting p-values were adjusted for multiple testing using the Benjamini–Hochberg method^90^.

### Retro-orbital injection

Twenty-five-month-old male C57BL/6 mice were anesthetized with isoflurane and received retro-orbital injections of AAV9-CAG-d2EGFP or AAV9-CAG-NKX2-5ΔHD at 1 × 10^13^ viral genomes per kg body weight. Mice were monitored until fully recovered from anesthesia before being returned to their home cages.

### Forelimb grip strength

Forelimb grip strength was measured using a grip strength meter (Columbus Instruments) following a protocol modified from a previously described method^91^. Briefly, mice were allowed to grasp a horizontal bar securely with their forelimbs and were gently pulled horizontally backward by the base of the tail until they released their grip. The meter recorded the peak force generated during each pull. Each trial comprised several gripping attempts to ensure secure engagement of both forepaws, and the highest valid peak force was recorded as the value for that trial. Nine attempts were performed for each mouse, with a 10-min intertrial interval during which the mice were returned to their home cages with ad libitum access to food and water. Both the mean and maximum grip-strength values across the nine attempts were used for analysis.

### Frailty index measurement

Mouse frailty was assessed using a previously described 31-item clinical frailty index^92^, with individual deficits scored according to published criteria^93^. Body weight and body surface temperature were scored on the basis of their deviations from the corresponding mean values in sex- and strain-matched young mice^92^. Forelimb grip strength was scored relative to the mean value of 60-day-old young mice, with strength ≥90%, 70–<90%, and <70% of the young mice assigned scores of 0, 0.5, and 1, respectively. All other deficits were scored as 0, 0.5, or 1 according to their severity based on instruction^93^. The individual deficit scores were summed and divided by the number of items assessed to generate an FI score ranging from 0 to 1, with higher scores indicating greater frailty.

**Fig. S1.**
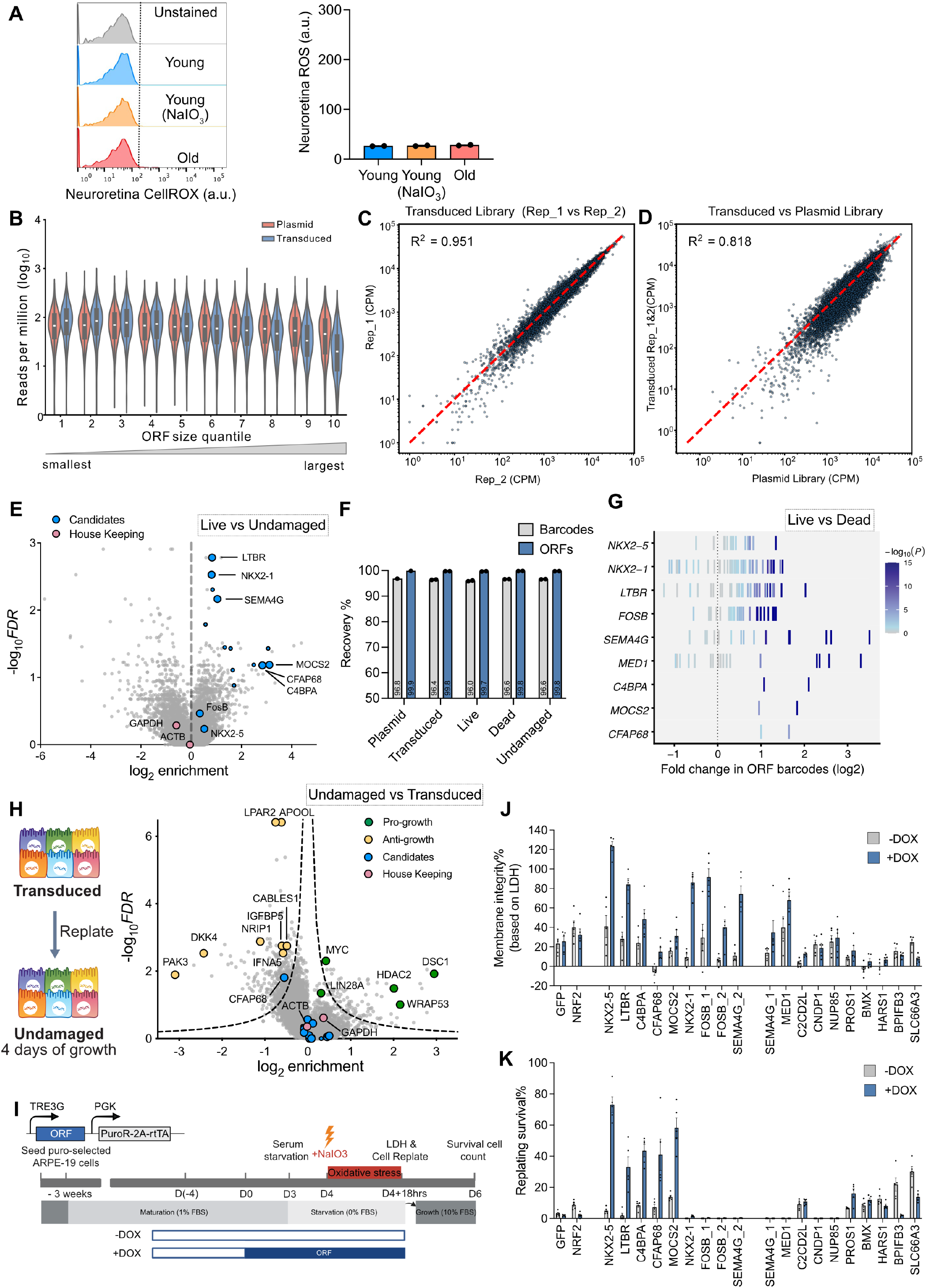
Oxidative-stress profiling, ORF screen quality control, and validation of protective candidates. **(A)** Representative flow cytometry histograms and quantification of CellROX signal in live neuroretina cells from NaIO_3_-treated young mice and aged mice compared with young controls (n= 2 in each group). **(B)** Distribution of reads pertaining to ORFs of varying sizes. ORFs were grouped into ten quantiles by size (Q1 = smallest, Q10 = largest; n ≈ 1,299 ORFs per quantile). Two conditions are shown: Plasmid library and the transduced library (an average of technical replicates). Reads are CPM-normalized and shown on a log_10_ scale. The box shows the 25th–75th percentiles with a line at the median, while whiskers extend to 1.5× the interquartile range. For ORFs that have isoforms, we assigned the largest isoform size. **(C)** Scatter plot comparing ORF abundance (CPM) between technical replicates of the transduced library, with each dot representing a single ORF, indicating robust reproducibility (R^2^ = 0.951). **(D)** Scatter plot comparing the average ORF abundance of the transduced library (two technical replicates) with the plasmid library. Each dot represents a single ORF, demonstrating high fidelity of library representation following lentiviral transduction (R^2^ = 0.818). **(E)** Volcano plot depicting differential enrichment of ORFs in live relative to undamaged cells. Each dot represents a single ORF. Gray dots indicate all screened ORFs, pink dots denote housekeeping genes, and blue dots highlight selected candidate ORFs subsequently tested in the arrayed validation assay. Smaller blue dots indicate false-positive hits, whereas larger blue dots indicate validated true-positive hits. **(F)** Bar chart showing uniform recovery (%) across sample conditions. Dots represent biological replicates (n = 2 for all conditions except the plasmid library, n = 1). Gray bars indicate barcode recovery, and blue bars indicate ORF recovery. **(G)** Barcode-level enrichment of selected candidate ORFs in the live versus dead comparison. Each tick represents an individual barcode, positioned by its log_2_ fold change (live versus dead) and colored by statistical significance (−log_10_(P)). Rows correspond to individual candidate ORFs, and the dotted line indicates no enrichment (log_2_FC = 0). **(H)** Volcano plot showing differential ORF enrichment in undamaged cells versus transduced cells after replating and expansion for 4 days. Each dot represents one ORF. **(I)** ARPE-19 cells with stably integrated doxycycline-inducible ORFs were matured in medium containing nicotinamide and 1% FBS^25^ for three weeks, then doxycycline (DOX) was added to induce ORF expression for independent survival assays. Validation of candidate protective factors in independent cell-survival assays. Cells expressing the indicated doxycycline-inducible ORF were cultured in the absence (−DOX) or presence (+DOX) of doxycycline and subjected to oxidative injury. **(J)** Membrane integrity was quantified from lactate dehydrogenase (LDH) release and expressed as the percentage of membrane integrity. Baseline membrane integrity (100%) was defined by LDH release from cells without DOX induction or NaIO_3_ treatment. Because starvation-medium exchange causes low background LDH release, values >100% indicate less membrane damage than baseline. **(K)** Long-term survival was assessed by replating cells after injury and quantifying the percentage of surviving cells. GFP and NRF2 served as controls. Bars represent group means with error bars, and individual dots represent biological replicates. Data are shown as mean ± s.e.m. Data are shown as mean ± s.e.m.

**Fig. S2.**
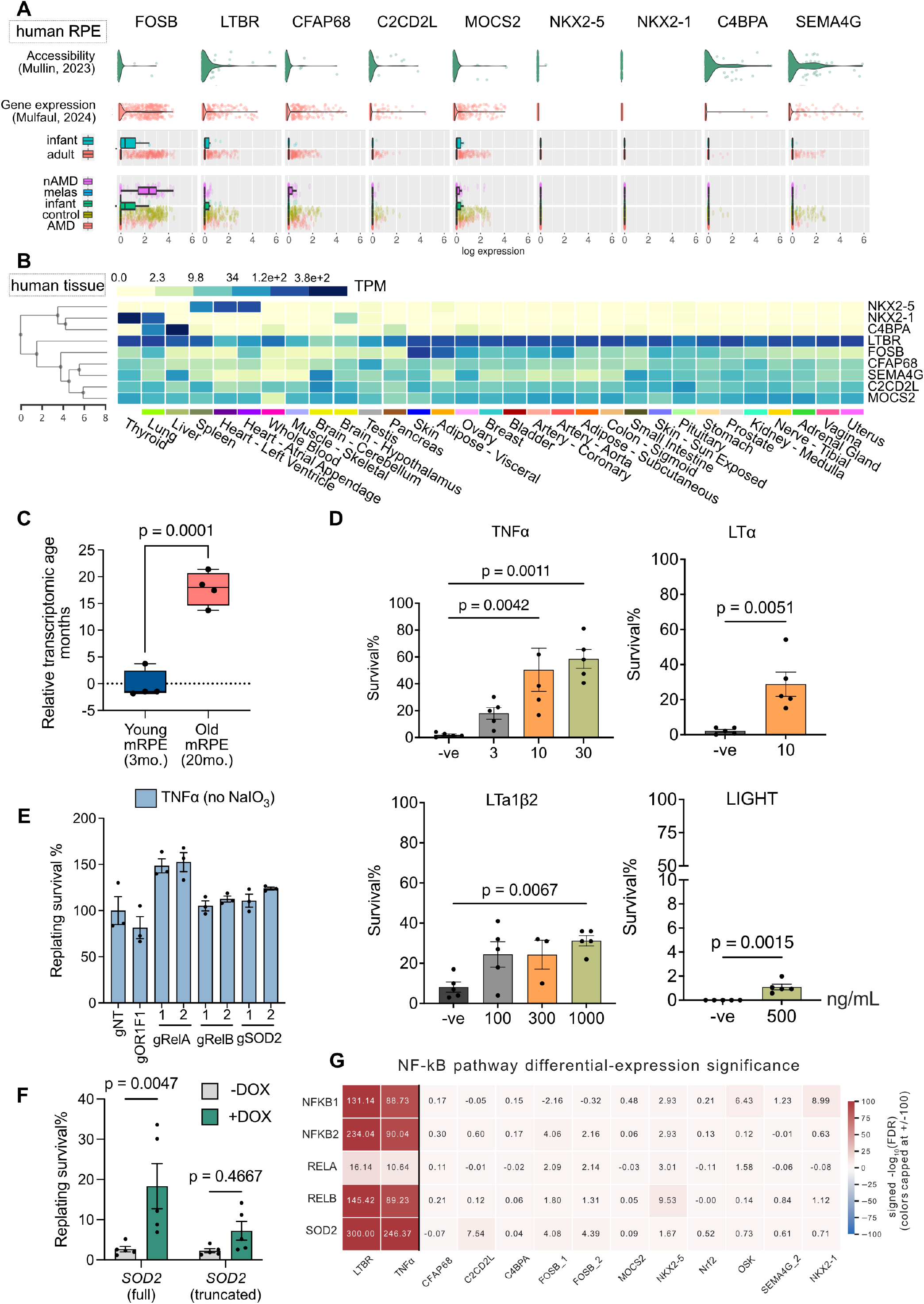
Candidate gene expression patterns and pathway-level transcriptional profiles. **(A)** RPE expression and chromatin accessibility profiles of selected candidate genes across human ocular single-cell and single-nucleus datasets^46,79^, including adult, aged, control, and AMD samples. **(B)** Tissue-specific expression heatmap of candidate genes based on normalized TPM values across human tissues. **(C)** Relative transcriptomic age prediction from RNA-seq data of young 3-month-old (n=4) and old 20-month-old (n=4) mouse RPE, calculated with the chronological rodent multi-tissue clock. **(D)** Survival of ARPE-19 cells treated with TNF-superfamily ligands at the indicated concentrations after NaIO_3_ challenge. **(E)** Replating survival of TNFα-treated ARPE-19 cells without oxidative stress following CRISPR-Cut targeting of RELA, RELB, or SOD2. Nontargeting (gNT) and *OR1F1*-targeting (gOR1F1) guides served as controls. Values were normalized to the mean of gNT, set to 100%. **(F)** Replating survival of ARPE-19 cells with doxycycline-inducible expression of full-length or truncated SOD2 after oxidative challenge. **(G)** Heatmap showing differential-expression significance for NF-κB pathway genes across candidate protective ORFs and controls. Values represent signed −log_10_(FDR). Data are shown as mean ± s.e.m. Data are shown as mean ± s.e.m.

**Fig. S3.**
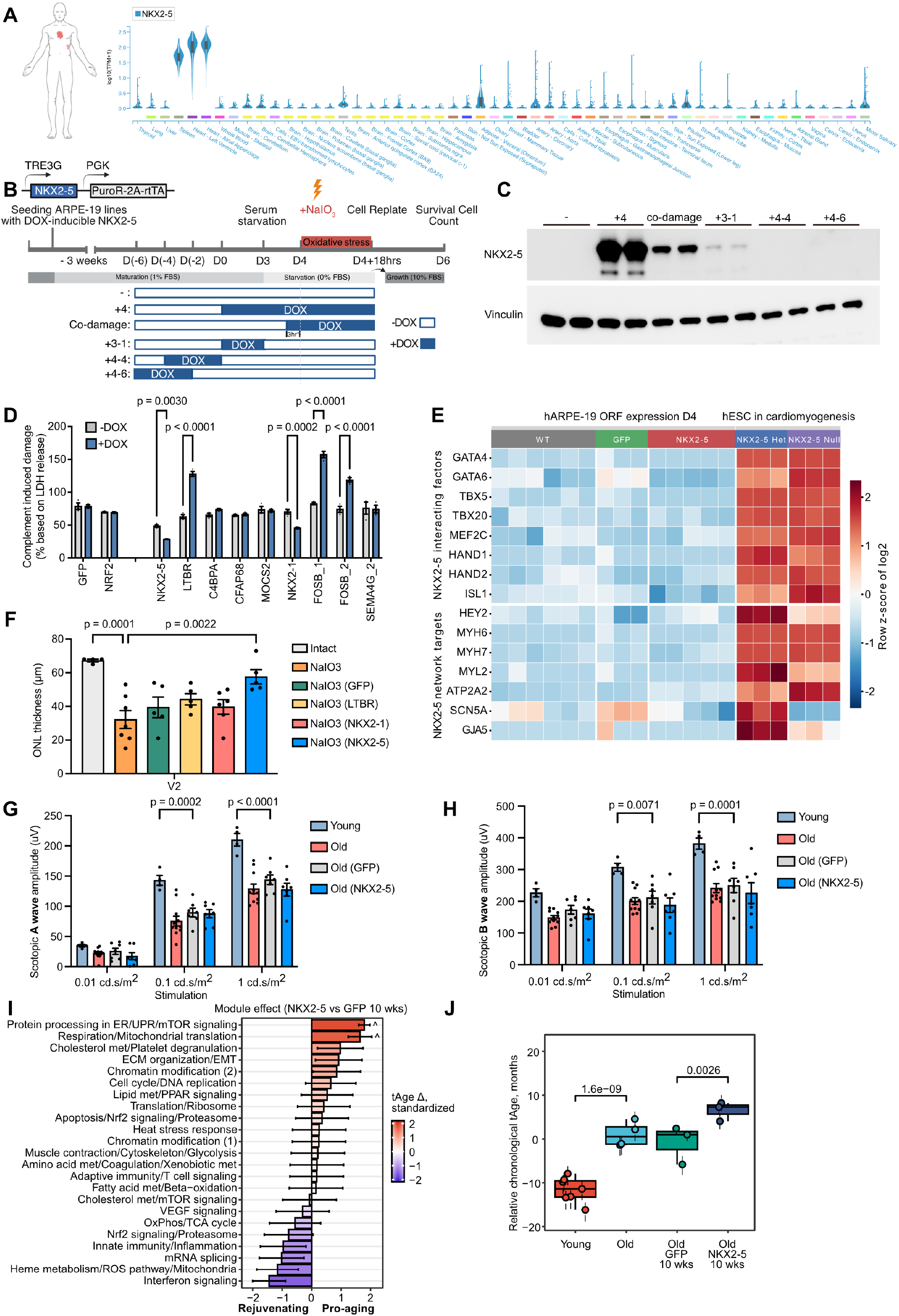
NKX2-5 expression, durability, retinal function, and transcriptomic effects. **(A)** Tissue-level expression profile of NKX2-5 across human tissues, showing restricted endogenous expression. **(B)** Experimental schematic for doxycycline-inducible NKX2-5 expression in ARPE-19 cells and timing of NaIO_3_ challenge, replating, and survival analysis. **(C)** Immunoblot analysis of NKX2-5 protein expression under the indicated doxycycline induction regimens. Vinculin was used as a loading control. **(D)** Complement-injury assay assessing the protective effect of ORF expression in ARPE-19 cells. Cells expressing GFP, NKX2-5, or the indicated genes were treated with 30% complement-containing human serum (CC-HS, Sigma, S1-LITER) for 7 days. Damage was normalized to LDH release from lysed cells in the undamaged control. Because LDH release during CC-HS treatment accumulates over 48 hours, normalized values can exceed 100%. **(E)** Analysis of cardiogenesis-associated gene networks in WT, GFP-, and NKX2-5-expressing RPE cells, with comparisons to RNA-seq samples of NKX2-5 Null and Het human embryonic stem cells in cardiomyogenesis (GSE89443). **(F)** Quantification of outer nuclear layer thickness in the untreated eye (n=4), NaIO_3_ injured eye (n=7), AAV-injected ventral region V2 after NaIO_3_ injury in eyes expressing GFP, LTBR, NKX2-1, or NKX2-5 (n=5, 5, 6 and 5 respectively). (**G, H**) Scotopic ERG responses from 17-month-old mice at 8 weeks after subretinal AAV-GFP or AAV-NKX2-5 injection. Scotopic a-wave amplitudes, reflecting predominantly photoreceptor activity, and b-wave amplitudes, reflecting downstream bipolar-cell and Müller glial responses, were measured across increasing flash intensities. Young mice (n=4), untreated old mice (n=11), and old mice 10 weeks after AAV-GFP (n=7) or AAV-NKX2-5 (n=7) treatment. (**I**) Module-specific transcriptomic age-clock analysis of aged RPE after AAV-NKX2-5 treatment compared with AAV-GFP controls at 10 weeks postinjection (GFP, n=3; NKX2-5, n=3). Red and blue indicate module-level shifts associated with increased and decreased transcriptomic age, respectively, calculated with the rodent multi-tissue chronological clocks. (**J**) Relative transcriptomic age estimates of young (n=7), old (n=4), AAV-GFP-treated old (n=3), and AAV-NKX2-5-treated old (n=3) RPE samples, calculated with the multi-tissue chronological clock. Data are shown as mean ± s.e.m.

**Fig. S4.**
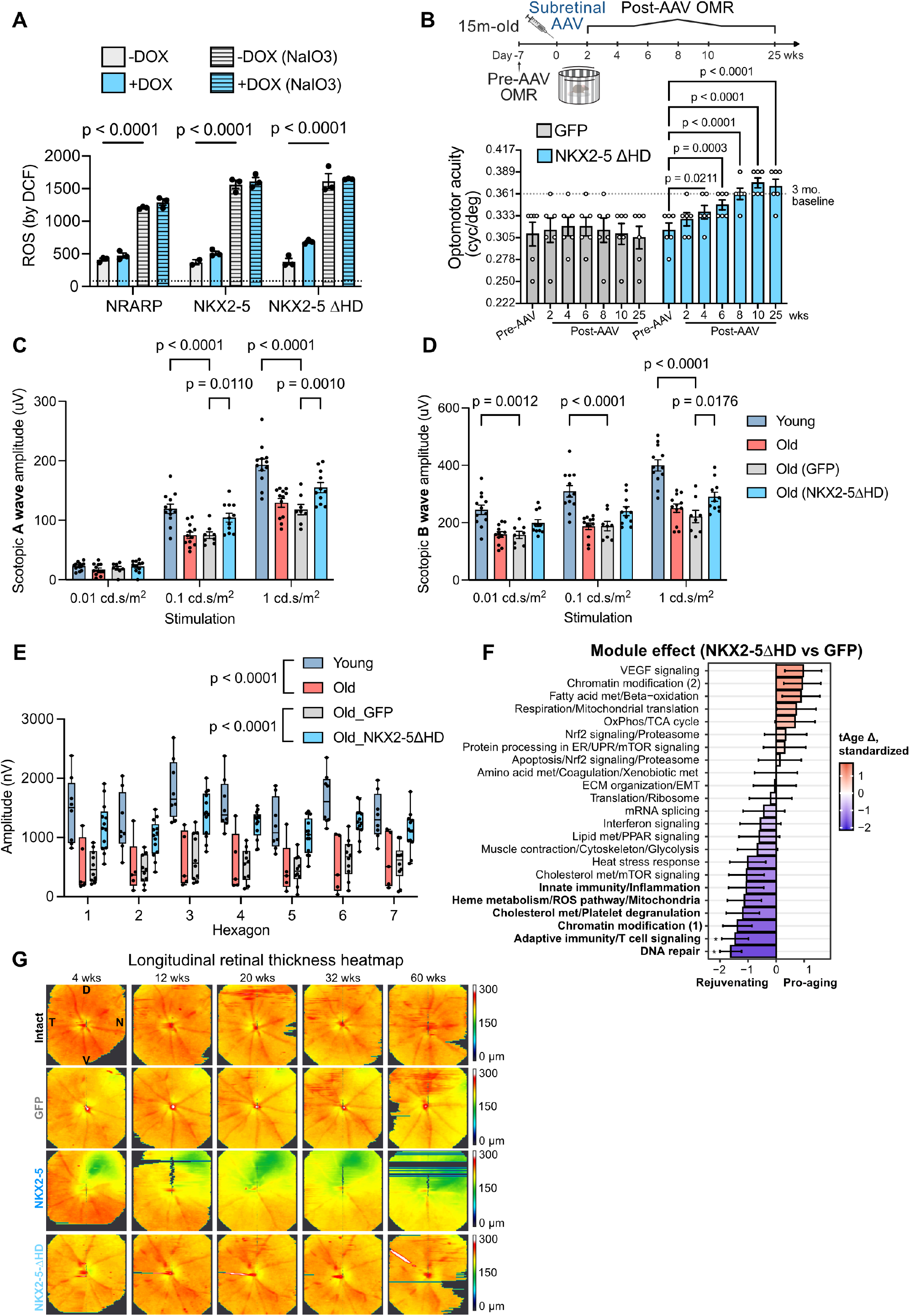
Characterization of NKX2-5ΔHD activity, aged retinal function, transcriptomic age, and long-term retinal structure. (**A**) ROS levels measured by DCF fluorescence in ARPE-19 cells expressing NRARP, full-length NKX2-5, or NKX2-5ΔHD, with or without doxycycline induction and NaIO_3_ challenge. Dotted line represents unstained values. (**B**) Longitudinal OMR measurements of visual acuity in aged mice after subretinal AAV-GFP (n=6) or AAV-NKX2-5ΔHD (n=6) injection, followed for up to 25 weeks post-AAV. The dotted line indicates the 3-month-old baseline reference. We could not perform ERG or OCT at the final time point because of high anesthesia-associated mortality in 21-month-old mice. Two-way repeated-measures ANOVA followed by Šídák multiple-comparison tests for comparisons with the pre-AAV baseline within each group. (**C, D**) Scotopic ERG responses from young mice (n=12), untreated old mice (n=12), and old mice treated with AAV-GFP (n=8) or AAV-NKX2-5ΔHD (n=11), two mice did not remain asleep for testing in each AAV group. Scotopic a-wave amplitudes (**C**) and b-wave amplitudes (**D**) were measured across increasing flash intensities. (**E**) Multifocal ERG responses across retinal hexagons in young mice (n=8), untreated old mice (n=5), and old mice treated with AAV-GFP (n=10) or AAV-NKX2-5ΔHD (n=13). (**F**) Module-specific transcriptomic age-clock analysis of old RPE treated with AAV-NKX2-5ΔHD (n=4) compared with AAV-GFP (n=4). Red and blue indicate module-level shifts associated with increased and decreased transcriptomic age, respectively, calculated with rodent multi-tissue chronological clocks. (**G**) Representative longitudinal total retinal thickness heatmaps from intact eyes and eyes treated with AAV-GFP, AAV-NKX2-5, or AAV-NKX2-5ΔHD, followed from 1 to 15 months post-AAV injection. Data are shown as mean ± s.e.m.

**Fig. S5.**
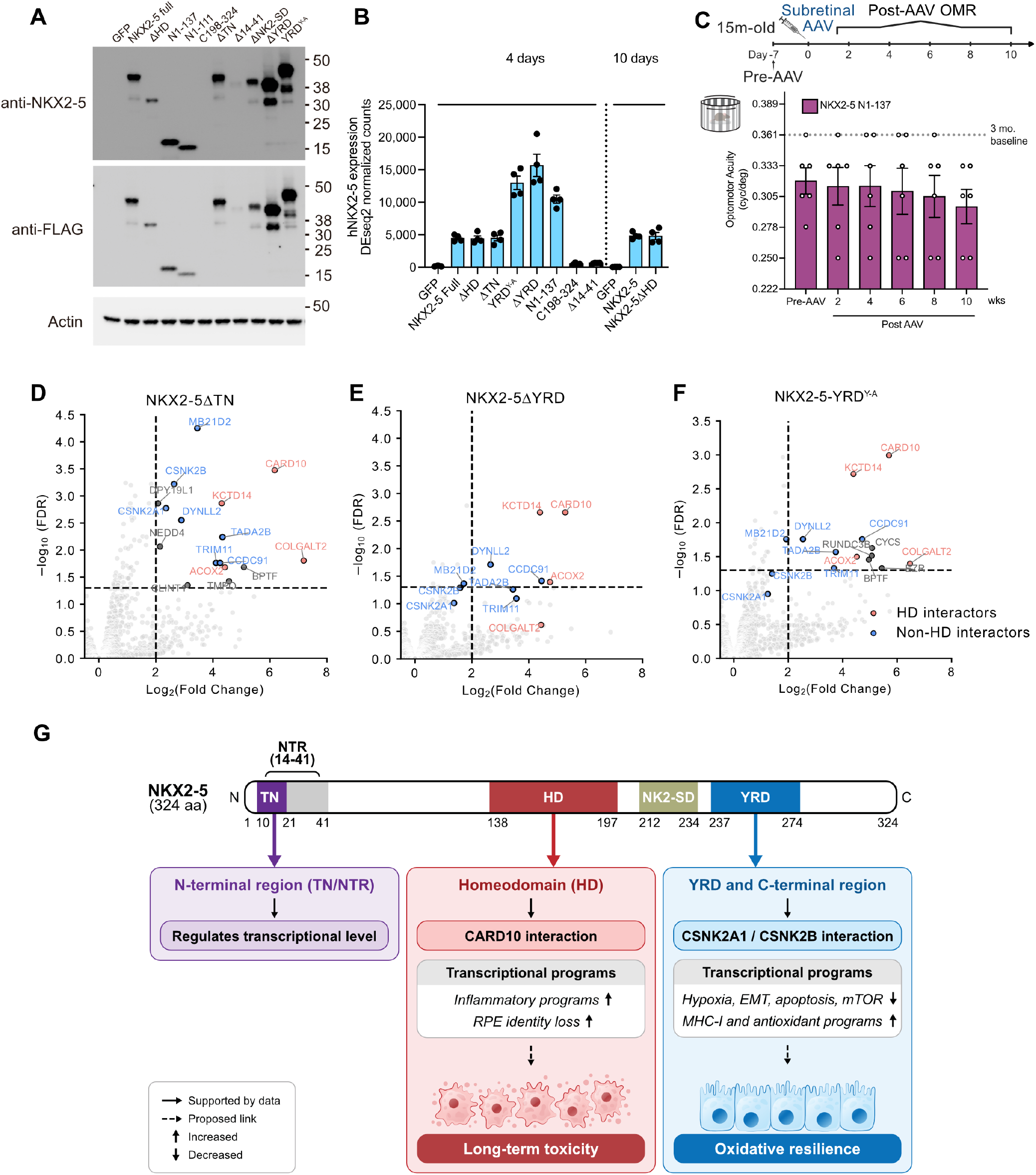
Domain-level analysis of NKX2-5 expression, pathway activity, interactions and function. (**A**) Immunoblot analysis of full-length NKX2-5 and indicated NKX2-5 mutants. NKX2-5 and FLAG antibodies were used to detect protein expression; actin served as a loading control. (**B**)Human *NKX2-5* transcript counts after expression of indicated NKX2-5 variants at 4 or 10 days. (**C**) Longitudinal OMR measurements after subretinal AAV expression of the NKX2-5 N-terminal fragment in aged mice (n=6). The dotted line indicates the 3-month-old baseline reference. (**D-F**) Co-IP–mass spectrometry analysis of ALFA-tagged NKX2-5ΔTN (**D**), NKX2-5ΔYRD (**E**) and NKX2-5-YRD^Y-A^ (**F**) binding partners. Volcano plots show enriched interactors. Previously identified HD-associated and non-HD-associated interactors are indicated. (**G**) Proposed model of NKX2-5 domain-associated activities. The N-terminal TN/NTR region is associated with regulation at the transcriptional level. The homeodomain (HD) is associated with CARD10 interaction, increased inflammatory programs and RPE identity loss, and long-term toxicity. The YRD/C-terminal region is associated with CSNK2A1/CSNK2B interactions, suppression of hypoxia, EMT, apoptosis, and mTOR programs, induction of MHC-I and antioxidant programs, and oxidative resilience. Data are shown as mean ± s.e.m.

**Fig. S6.**
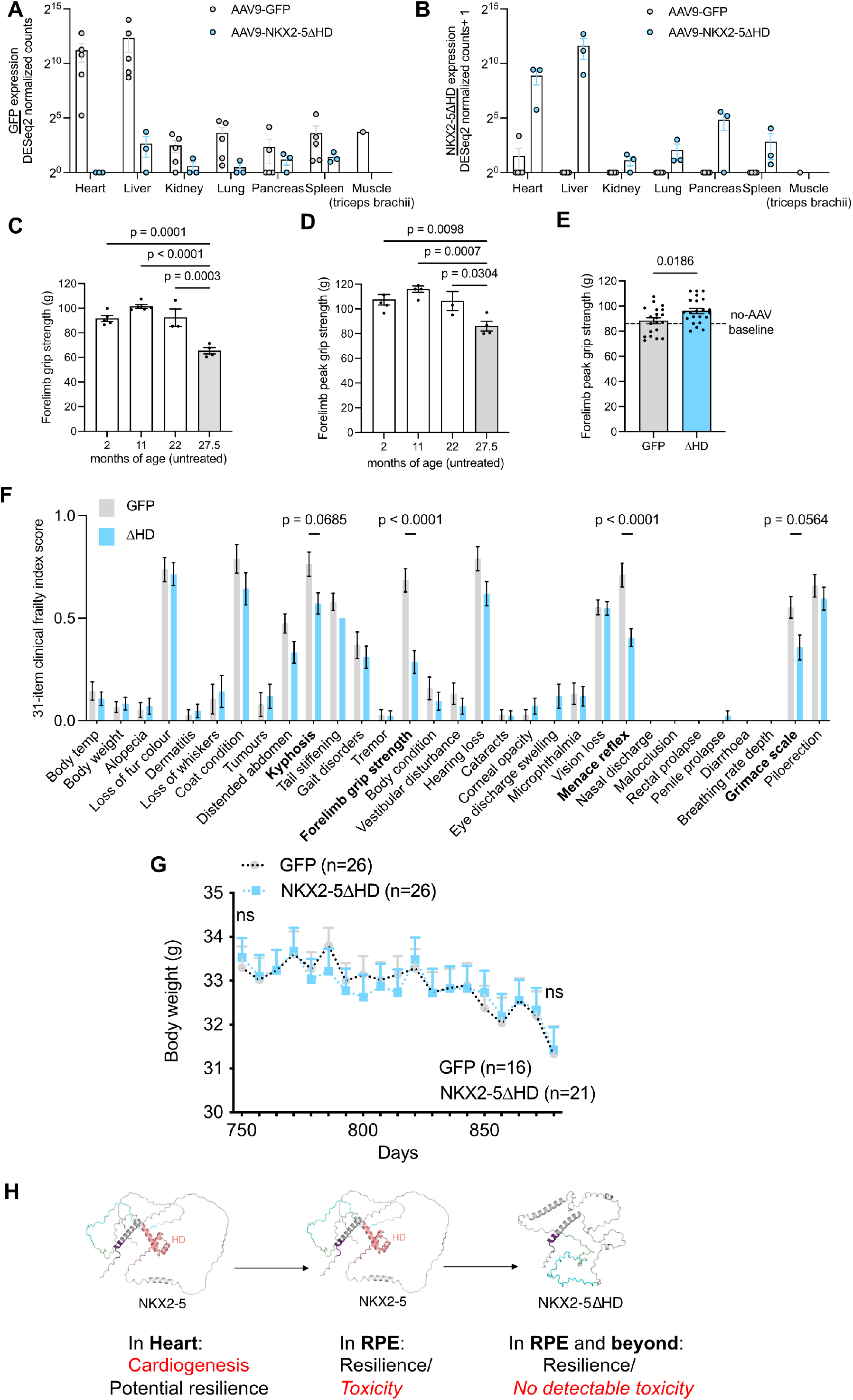
Systemic delivery of NKX2-5ΔHD improves grip strength and reduces frailty in aged male mice. **(A)** GFP transcript abundance across tissues following systemic AAV9 delivery. GFP expression was quantified in heart, liver, kidney, lung, pancreas, spleen, and triceps brachii muscle from mice treated with AAV9-GFP (n=5 except n=1 for muscle) or AAV9-hNKX2-5ΔHD (n=3). Expression is shown as log transformed DESeq2-normalized counts. Each dot represents an individual animal; bars show mean ± SEM. **(B)** hNKX2-5ΔHD transcript abundance across tissues following systemic AAV9 delivery. NKX2-5ΔHD expression was quantified in heart, liver, kidney, lung, pancreas, and spleen from mice treated with AAV9-GFP (n=5 except n=1 for muscle) or AAV9-hNKX2-5ΔHD (n=3). Expression is shown as log transformed DESeq2-normalized counts. Each dot represents an individual animal; bars show mean ± SEM. **(C)** Forelimb grip strength of untreated male C57BL/6 mice at the indicated ages. Grip strength was calculated as the average of repeated attempts. **(D)** Peak forelimb grip strength of untreated male C57BL/6 mice at the indicated ages (n=5, 5, 3 and 4, respectively). Peak grip strength was defined as the maximum value across repeated attempts. **(E)** Peak forelimb grip strength of AAV9-GFP- and AAV9-NKX2-5ΔHD-treated male mice at 27.5 months of age, 80 days after AAV injection (GFP, n = 20; NKX2-5ΔHD, n = 22). Peak grip strength was defined as the maximum value across the nine attempts. **(F)** Itemized 31-item clinical frailty index scores in aged mice treated with AAV9-GFP or AAV9-hNKX2-5ΔHD (GFP, n = 19; NKX2-5ΔHD, n = 21). Bars show mean scores for each frailty item, with lower scores indicating reduced severity of frailty. Examiners were masked to the group allocation. P values were calculated using two-way ANOVA, P <0.1 are labelled. **(G)** Longitudinal body weight of aged mice treated with AAV9-GFP or AAV9-hNKX2-5ΔHD, starting with n = 26 in each group. P values were calculated using two-way ANOVA. **(H)** Schematic of engineering NKX2-5 to separate resilience from toxicity in RPE. Native NKX2-5 contains a homeodomain (HD) that supports its canonical role in cardiogenesis and tissue resilience in the heart. When ectopically expressed in RPE, full-length NKX2-5 promotes oxidative resilience but is associated with toxicity. Deletion of the homeodomain generates NKX2-5ΔHD, which retains the resilience-promoting activity with no detectable retinal toxicity under the conditions tested. Data are shown as mean ± s.e.m.

