## Supplementary Discussion for "A genome-scale ORF screen for resilience factors that protect the RPE and restore vision in aged mice"

Our lab and others have shown that mice display an age-related loss of visual function as measured by a significant reduction in OMR and ERG[1–3]. Our current results showed that AAV2-NKX2-5 treatment of RPE in aging mice significantly reversed their age-induced decline in visual acuity as assessed by OMR (**Fig.3I**) but no ERG improvements were observed (**Fig.3J, and Fig.S3, G and H**). Such discordance between behavioral and electrophysiological measurements has been reported across multiple models of retinal degeneration[4–6]. Notably, in our previous AAV-OSK studies, prolonged OSK expression similarly improved OMR while eventually reducing ERG responses[6], providing a directly relevant precedent for divergence between these functional readouts after AAV-mediated intervention. A related dissociation between temporal contrast sensitivity and ERG has also been reported in a mouse model of retinitis pigmentosa, in which behavioral visual performance improved despite declining ERG responses[7].

For example, across eight transgenic rat models of retinal degeneration, OKT and ERG showed markedly different trajectories depending on the rate of degeneration: slowly degenerating models displayed rapid declines in OKT thresholds that were not matched by ERG changes, whereas rapidly degenerating models showed comparatively slow OKT decline despite substantial electrophysiological deterioration[4]. Similarly, in a mouse model of paraquat-induced retinal oxidative stress, ERG a-wave and OMR measurements correlated poorly at low levels of retinal injury, with their correlation increasing only as oxidative damage became more severe[5]. Together, these studies indicate that behavioral visual responses and full-field ERG interrogate overlapping but distinct aspects of retinal function.

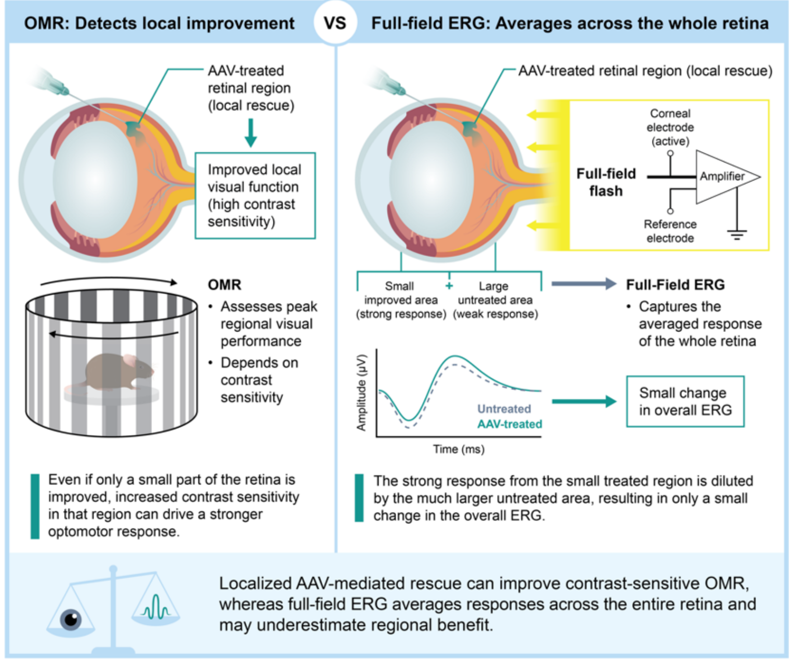

**Localized AAV rescue can diverge from full-field responses**

One explanation particularly relevant to subretinal AAV delivery is the spatial difference between these measurements. Subretinal injection transduces a localized retinal region, and functional restoration within this area may be sufficient to improve visually guided behavior. In contrast, full-field ERG measures the summed electrical response of the entire retina and can therefore dilute localized improvements, a recognized limitation when evaluating focal retinal interventions [8].

Systematically testing this relationship across additional AAV interventions is beyond the scope of the present study. Nevertheless, these observations highlight the value of combining behavioral measurements, full-field ERG, and spatially resolved electrophysiology when evaluating localized AAV-based retinal therapies. Importantly, AAV-NKX2-5ΔHD showed consistent benefits across multiple functional readouts, supporting a safer profile than full-length NKX2-5.
